# Backbone Hydrogen Bonding as a Determinant of Condensate Material States

**DOI:** 10.64898/2025.12.21.695814

**Authors:** Yi Zhang, Ramesh Prasad, Huan-Xiang Zhou

**Affiliations:** Department of Chemistry, University of Illinois Chicago, Chicago, IL; Department of Physics, University of Illinois Chicago, Chicago, IL

## Abstract

Condensate material states ranging from liquid droplets to gels have direct functional consequences, but their molecular determinants, especially for gels, are poorly understood. Here we combined optical microscopy, NMR spectroscopy, and molecular dynamics (MD) simulations to test the hypothesis that backbone hydrogen bonding is a determinant for the material states of XYssYX tetrapeptides. IAssAI forms droplets but AIssIA forms gels; MD simulations show a higher level of hydrogen bonds in AIssIA condensates than in IAssAI condensates. Addition of a low-hydrogen-bonding peptide, AAssAA, converts AIssIA gels into droplets. The same effect is achieved by methylating the backbone amides of AIssIA to block hydrogen bonding. ^1^H-^1^H nuclear Overhauser effect spectroscopy, coupled with MD simulations, reveals strong, backbone hydrogen bonding-buttressed molecular networks in AIssIA gels. These results demonstrate that, in addition to sidechain-sidechain interaction strength, backbone hydrogen bonding, by promoting directional growth, is a determinant of condensate material states.

## Introduction

After formation via phase separation, biomolecular condensates exhibit a variety of material states ^1–6^. Phase separation can occur spontaneously via spinodal decomposition, which starts with concentration fluctuations leading to interconnected domains ^7^. The most common material state is liquid droplets (Figure 1a), inside which intermolecular interactions form and break; droplets have a spherical shape and can fuse, both driven by interfacial tension. P granules, Cajal bodies, nucleoli, and stress granules possess these liquid-like properties ^8–11^. The second material state is reversible aggregates (Figure 1b), characterized by overly strong intermolecular interactions ^12^, which prevent the completion of phase separation and produce particles with irregular shapes. The third material state is gels (Figure 1c), which emerge from arrested spinodal decomposition ^13^ and can grow into system-spanning networks. By deep quenching into the spinodal region, the N-terminal low-complexity region (FUS_N_; residues 1-214) fused with a light-activatable domain (upon light activation) ^14^ and the peptide carboxybenzyl-capped di-phenylalanine (upon temperature decrease) ^15^ formed gels. The FG-rich domain of the yeast nucleoporin Nsp1p also formed gels; gelation is required for nuclear pore complex function ^16^. P granules contain a peripheral gel phase that provides a stabilizing scaffold and helps with their localization in the posterior cytoplasm ^17^. Cytoplasmic-abundant heat-soluble (CAHS) proteins in tardigrades undergo a gel transition to increase cell stiffness during dehydration ^18^. Gels can be very porous and filled with a high level of water and thus are often referred to as hydrogels ^16, 18–22^. A fourth material state, amorphous dense liquids (Figure 1d), has been reported recently ^23^ to occur when overly weak intermolecular interactions prevent further condensation shortly after spinodal decomposition starts. Transient clusters formed by the Mediator coactivator and RNA polymerase II are possible examples of amorphous dense liquids ^24^. Lastly, many biomolecular condensates age into irreversible aggregates ^25^.

**Figure 1.**
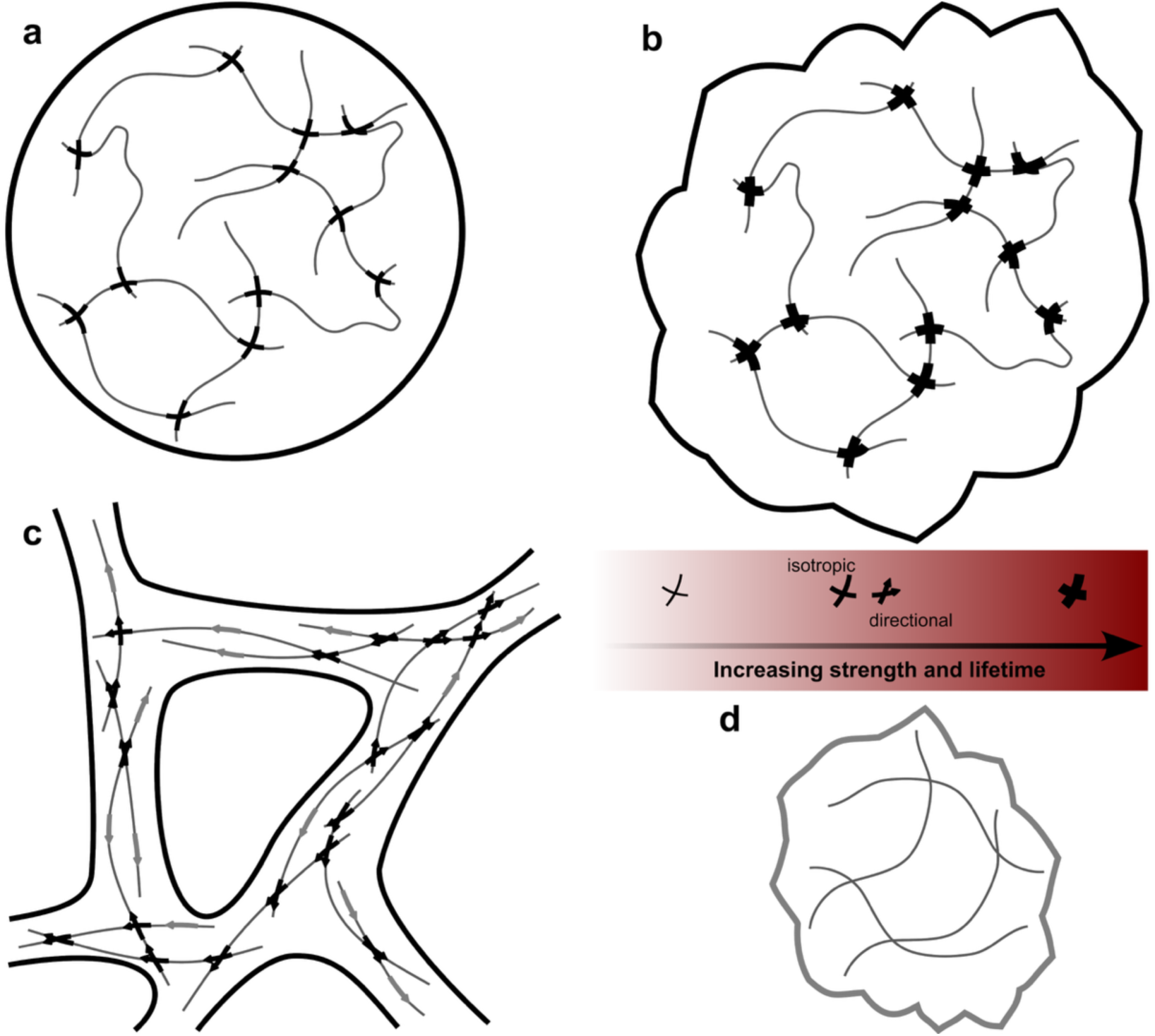
Intermolecular networks inside four material states of biomolecular condensates. (a) Droplets occur after complete phase separation. Moderately strong interactions allow intermolecular crosslinks to form, break, and reform quickly, giving rise to liquidity. The remaining three states result from the termination of phase separation at various stages of spinodal decomposition (dense regions arising from concentration fluctuations; interconnected domains upon further condensation; breakage of interdomain necks; and domains morphing into a spherical shape). (b) Reversible aggregates occur when phase separation is terminated after the breakage of interdomain necks; overly strong interactions prevent further rearrangement inside domains. (c) Gels occur when phase separation is terminated when domains are still interconnected; directional interactions prevent breakage of interdomain necks. (d) Amorphous dense liquids occur when phase separation is terminated at a very early stage, as overly weak interactions prevent further condensation.

Among the four types of “fresh” (i.e., non-aged) condensates, amorphous dense liquids, liquid droplets, and reversible aggregates follow the order of increasing intermolecular attraction (Figure 1). Correspondingly, the dynamics of these three types of condensates follow a decreasing order, as reported by slowing fluorescence recovery after photobleaching (FRAP), slowing fusion, and increasing viscosity ^23, 26–28^. In comparison, gels are much less understood. For example, FRAP can be very slow in gel networks and gel samples lose fluidity; by these properties gels may be viewed as more solid-like than reversible aggregates, and yet intermolecular attraction may not be stronger in gels than in aggregates ^23^. For FUS, Kato et al. ^19^ found that the disordered FUS_N_ was necessary and sufficient for gel formation at high concentrations. Moreover, they identified a highly enriched motif [G/S]Y[G/S]; the ability to form gels attenuated when an increasing number of middle Y residues were mutated to S residues. Likewise, the FG-rich domain of the yeast nucleoporin Nsp1 is very likely to be disordered, and mutation of all its 55 F residues into S residues abolished gel formation ^16^. The disordered histidine-rich beak protein HBP-1 and its N-terminal deletion mutants formed droplets but its C-terminal deletion mutants formed hydrogels; designed peptides using repetitive units from HBP-1 suggest that a GAGFA unit favors gel formation ^21^. On the other hand, for Pab1, intrinsically disordered regions (IDRs) were dispensable for gel formation but modulated phase-separation propensity ^20^. Similarly, the folded domain at the C-terminus of MEG-3, the protein that forms the gel phase in P granules, is required, as its deletion results in the loss of MEG-3’s ability to form a distinct phase ^17^. The gelation of CAHS proteins required coiled-coil regions, as deletion of and L-to-P mutations in these regions abrogated gel transition ^18^. LplA, a structured protein, readily underwent a gel transition upon a temperature rise from 4 °C to 25 °C ^22^. For SPOP-DAXX cocondensates, it was proposed that the driving force for gel formation is the entropy of binding-site occupation by DAXX on SPOP ^29^. In coarse-grained simulations, introducing intermolecular β-sheet formation led to gelation ^30^. These disparate observations show that we are still at an early stage of characterizing molecular determinants of gels.

Short peptides are very useful in characterizing and dissecting the drivers of gel formation by proteins. For example, a shallow quench (temperature drop from 80 °C to 60 °C) of carboxybenzyl-capped di-phenylalanine produced liquid droplets but a deep quench (temperature drop from 80 °C to 20 °C) produced gels ^15^. A similar behavior was observed on FUS_N_ and, indeed, expected for systems that form gels through arrested spinodal decomposition. C-terminal amidated tripeptides consisting of D, F, and Y exhibited a variety of phase behaviors ^31^. FDY and YDF did not phase separate, FYD formed aggregates, whereas YFD, DFY, and DYF formed gels. The DFY gel grew from crystalline fibers. X-ray diffraction revealed fiber stabilization by hydrogen bonds between backbone amides (β-sheet-like) and between Y hydroxyls, salt bridges between D carboxyls and N-terminal amines, and π-π stacking of the aromatic rings. Interestingly, when Y-Y hydrogen bonds were eliminated by Y oxidation, gels were converted to droplets. Fluorenylmethyloxycarbonyl (Fmoc)-capped KKYY formed gels at neutral pH; upon Y phosphorylation, the peptide formed gels at acidic pH ^32^. Based on aggregation propensities (defined by reduction in solvent accessible surface area) in coarse-grained simulations, a machine-learning method was developed to select 165 tetrapeptides, of which 100 formed gels. Among the latter peptides, F had the highest probabilities of being at the fourth and third positions (37% and 29% respectively), whereas V had the highest probability (26%) at the second position and L had the highest probability (23%) at the first position ^33^. Abbas et al. ^34^ studied tetrapeptides of the form YFzFY, where z is short for NH-CH_2_-CH_2_-Z-CH_2_-CH_2_-NH (see Figure 2a for Z = S-S). YFoFY and YFsFY formed droplets but YFcFY and YFssFY formed gels. We recently reported that FFssFF, LLssLL, and MMssMM formed droplets whereas IIssII and VVssVV formed gels ^23^. All-atom molecular dynamics (MD) simulations suggested that the gel-forming peptides had higher propensities of forming backbone hydrogen bonds. This suggestion was validated by the effects of urea addition: whereas 3 M urea had only a modest effect on FFssFF droplet formation, it dissolved IIssII gels and converted VVssVV gels into amorphous dense liquids.

**Figure 2.**
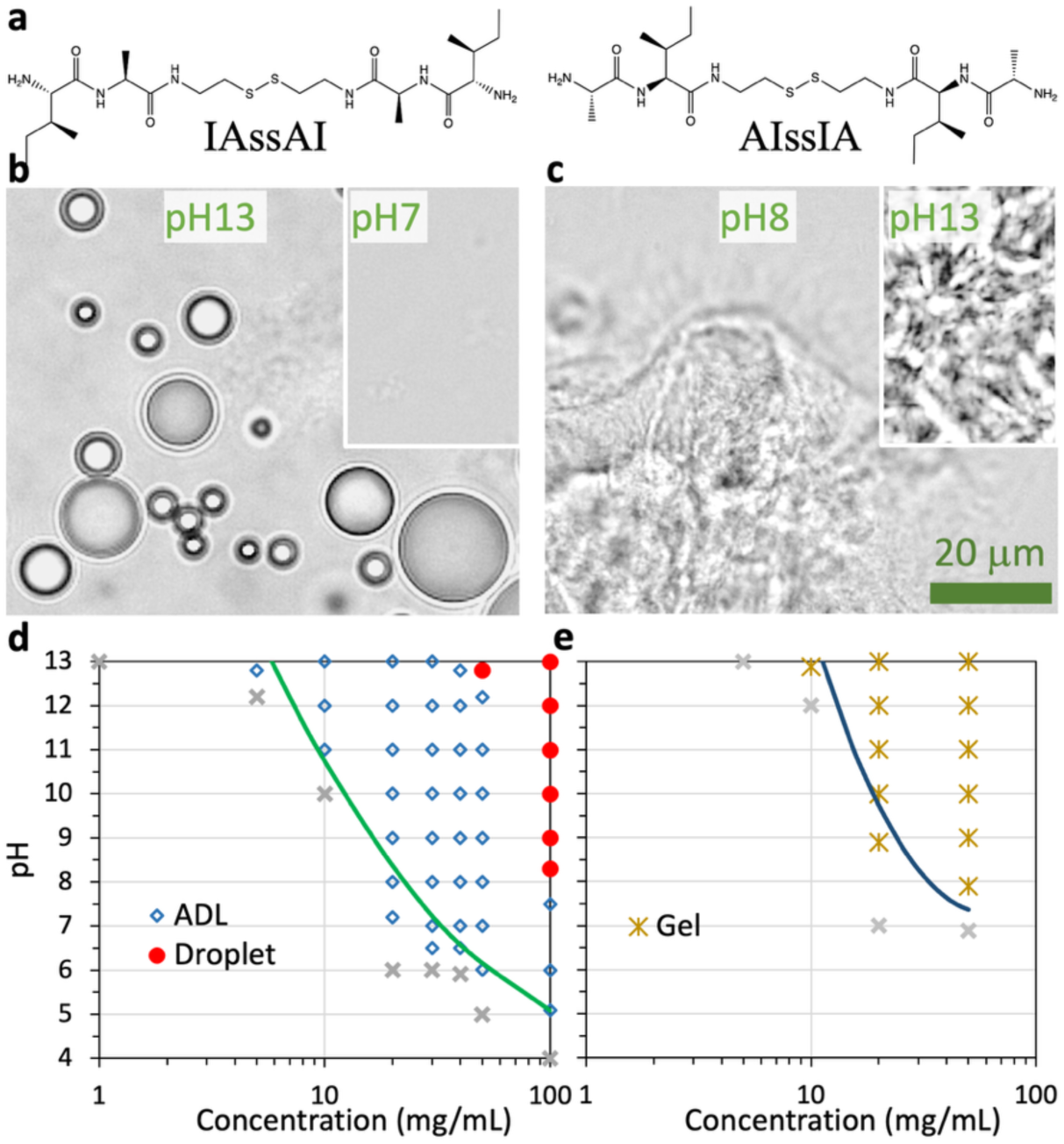
AIssIA and IAssAI form condensates with different material states. (a) Chemical structures of the two peptides. The terminal amines are shown in the charge-neutral form, appropriate for high pH; at low pH, each amine can bond with an additional proton, resulting in a +1 charge. (b) Brightfield images of IAssAI condensates at 100 mg/mL in Milli-Q water, showing droplets at pH 13 but amorphous dense liquids at pH 7. (c) Brightfield images of AIssIA condensates at 100 mg/mL in Milli-Q water, showing gels at both pH 8 and 13. (d, e) Phase diagram of IAssAI (panel d) and AIssIA (panel e) in 50 mM imidazole buffer.

The roles of hydrogen bonding in phase separation have been studied by various methods. In some cases, the fibers making up gel networks are crystalline; X-ray diffraction can reveal molecular arrangements, including backbone and sidechain hydrogen bonds ^31^. Another structural method, solution NMR spectroscopy, has indicated that hydrogen bonding between a Y and an H residue was the initial step in the phase separation of an HBP-1 fragment ^21^. The second approach is to introduce modifications to either eliminate or add hydrogen bonds. For example, oxidation of Y eliminates its ability to form sidechain hydrogen bonds ^31^. Similarly, methylation of amide nitrogen prevents backbone hydrogen bonding ^35, 36^. On the other hand, hydroxylation of Y introduces a second hydroxyl on the phenol ring, and the increased hydrogen bonding ability is essential for the phase separation of an adhesive peptide derived from the mussel foot protein Mfp-5 ^37^. The third approach is to use an additive like urea to perturb hydrogen bonding strength, as demonstrated in several studies ^23, 37–39^. Complementing these experimental approaches, all-atom MD simulations can enumerate all intermolecular interactions, including hydrogen bonds, that drive phase separation ^23^.

Our previous observation that LLssLL formed droplets but IIssII formed gels ^23^, despite the chemical similarity between L and I, underscores the mystery surrounding gel formation. The mystery deepened when we discovered here that IAssAI formed droplets but AIssIA formed gels, implicating the order of amino acids along the sequence as an important factor for condensate material states. We thus set out to test the hypothesis that backbone hydrogen bonding is a determinant of condensate material states. Specifically, we tested whether backbone hydrogen bonding promotes gel formation of AIssIA, by combining optical microscopy, NMR spectroscopy, and MD simulations. MD simulations indeed show a higher level of hydrogen bonds in AIssIA condensates than in IAssAI condensates. In concordance, 2 M urea dissolves AIssIA gels but not IAssAI droplets. Moreover, addition of a low-hydrogen-bonding peptide, AAssAA, and methylation of backbone amides convert AIssIA gels into droplets or amorphous dense liquids. Importantly, ^1^H-^1^H nuclear Overhauser effect spectroscopy (NOESY) coupled with MD simulations reveals strong, backbone hydrogen bonding-buttressed molecular networks in AIssIA gels, but more transient intermolecular association in IAssAI droplets. These results demonstrate that, in addition to sidechain-sidechain interaction strength, backbone hydrogen bonding, by promoting directional growth, is a determinant of condensate material states.

## Results

We used a pH-responsive tetrapeptides of the form XYssYX to elucidate the molecular determinants of condensate material states. At pH 2, the terminal amines each carry a positive charge upon protonation; the resulting charge-charge repulsion prevents phase separation. A pH rise (via adding NaOH) triggers phase separation to form condensates with various material states. By adding urea, mixing with a low-hydrogen-bonding peptide, or introducing N-methylation, we show that AIssIA gels can be dissolved or converted into droplets or amorphous dense liquids. Lastly, we combine ^1^H-^1^H NOESY and MD simulations to characterize the intermolecular interactions, in particular backbone hydrogen bonding, that differentially stabilize gels and other types of condensates.

### IAssAI forms droplets but AIssIA forms gels, implicating the order of amino acids along the sequence as an important factor for material states

Previously we have shown that, under phase-separation conditions, LLssLL forms either droplets (at pH > ∼7) or amorphous dense liquids (at lower pH), whereas IIssII forms only gels ^23^. With two different amino acids, A and I, two different heteropeptides can be generated: IAssAI and AIssIA (Figure 2a). Remarkably, the two peptides condense into different material states. 100 mg/mL IAssAI forms droplets at pH 13 and amorphous dense liquids at pH 7 (Figure 2b), whereas 100 mg/mL AIssIA forms gels at both pH 8 and pH 13 (Figure 2c). We captured the initial formation and growth of gel networks in a video (Video S1), where two drops of 5-M NaOH were pipetted into a 1-mL AIssIA sample at 10 mg/mL (just inside the gel region; see next). Gels immediately form in the wake of the falling NaOH drops; over time, the gels grow laterally.

We determined the phase diagrams of the two peptides over a range of pH and concentrations. For IAssAI, amorphous dense liquids occupy most of the phase-separation region; droplets only take up a corner at high pH and high concentration (Figure 2d). In contrast, for AIssIA, gels occupy the entire phase-separation region (Figure 2e). Thus qualitatively, the phase diagrams of IAssAI and AIssIA are similar to those of LLssLL and IIssII, respectively. It is not clear what is in common between LLssLL and IAssAI and between IIssII and AIssIA; nor is it clear how a change in the order of the two amino acids, A and I, along the sequence dictates the condensate material state.

### MD simulations reveal backbone hydrogen bonding as a determinant of material states

To address these questions, we turned to MD simulations. We started the simulations with 64 chains of each peptide in the high-pH form, uniformly distributed in a cubic box, and relied on spinodal decomposition for spontaneous phase separation ^40^, similar to the experimental conditions. In agreement with the experimental observations, upon phase separation, the IAssAI system forms a flat slab, which is equivalent to spherical droplets if not for the periodic boundary condition imposed in the simulations, whereas the AIssIA system forms gel networks (Figure 3a, b). The gel networks are stabilized not only by side-chain hydrophobic clusters but also by backbone hydrogen bonds (magenta lines in Figure 3b and inset). The average number of hydrogen bonds formed by one half of each AIssIA chain is 0.46, similar to the corresponding values of 0.55 and 0.44 for the previously studied gel-forming peptides IIssII and VVssVV, respectively ^23^ (Figure 3c). In comparison, the corresponding value, 0.37, for the droplet-forming IAssAI is much lower and close to the counterpart, 0.36, for the droplet-forming LLssLL ^23^. The value for AAssAA, which forms amorphous dense liquids, is even lower, at 0.29 ^23^.

**Figure 3.**
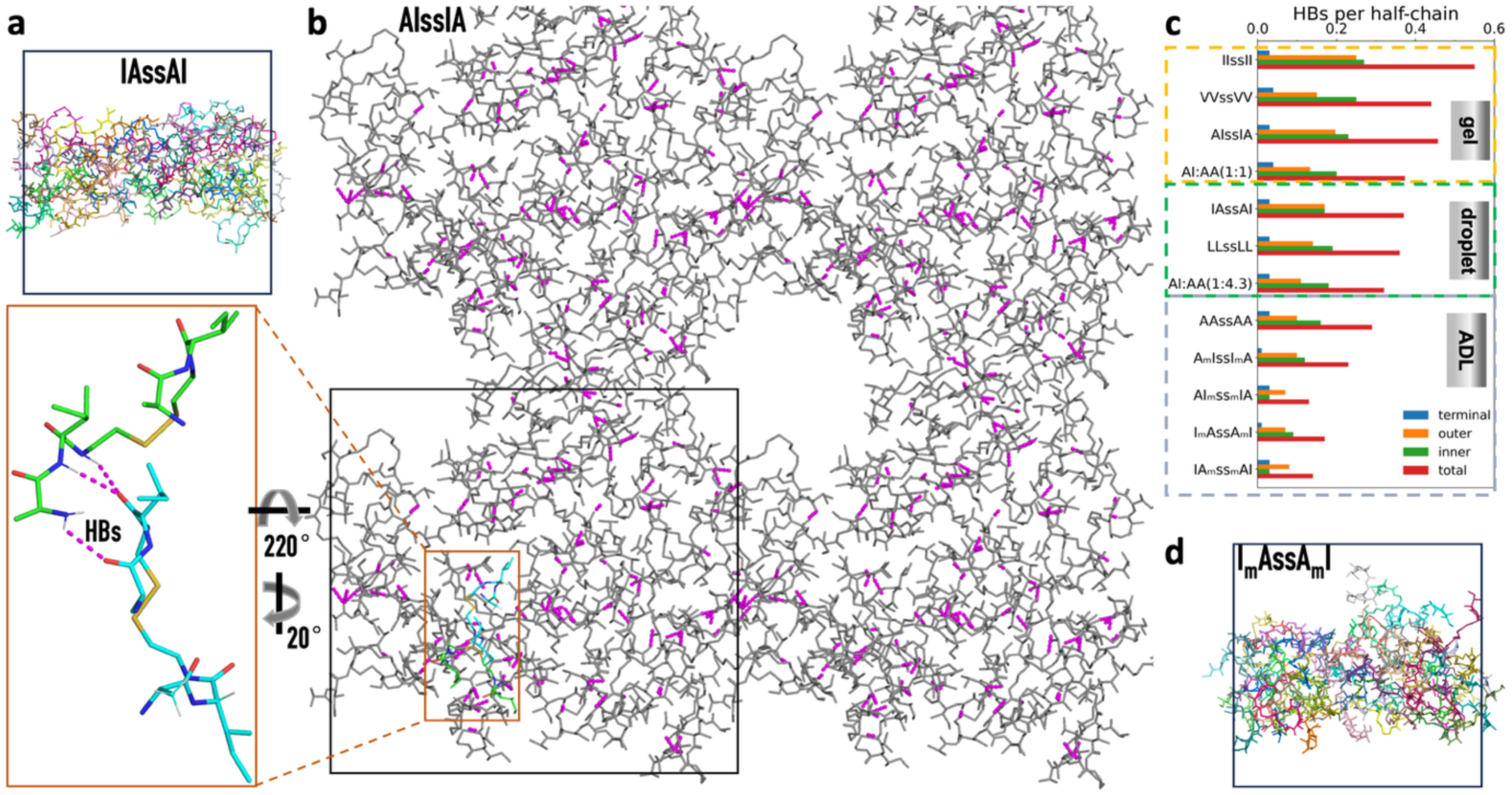
All-atom MD simulations reveal backbone hydrogen bonding as a determinant of material states. (a) A flat slab formed by IAssAI chains, corresponding to droplets. (b) Gel networks formed by AIssIA chains. Peptide chains in the primary unit cell (indicated by a black box) and three image cells are displayed, with interchain hydrogen bonds shown as magenta lines. A zoomed view highlights hydrogen bonds (HBs) between two chains. (c) Average number of hydrogen bonds formed by one half of each peptide chain. This number is further broken into contributions by the terminal amine, outer amide, and inner amide. Results for IIssII, VVssVV, LLssLL, and AAssAA were reported previously ^23^. ADL: amorphous dense liquid. (d) A slab with rough surfaces formed by outer N-methylated IAssAI, corresponding to amorphous dense liquids.

### 2 M urea dissolves AIssIA gels by disrupting backbone hydrogen bonding

To verify that hydrogen bonding is indeed a driver for the difference in material states between IAssAI and AIssIA, we added urea to the samples. Urea is thought to disrupt hydrogen bonds while preserving hydrophobic interactions. We chose peptide concentrations just inside the phase boundaries for IAssAI droplets and AIssIA gels (Figure 2d, e), i.e., 75 mg/mL IAssAI and 7.5 mg/mL AIssIA at pH 13. With 2 M urea, IAssAI droplets are still observed, though the number is reduced (Figure 4a). In contrast, 2 M urea completely dissolves AIssIA gels (Figure 4b). The differential response to urea addition suggests that AIssIA exhibits a higher propensity for hydrogen bonding than IAssAI, consistent with the MD simulation results (Figure 3a-c). The MD and urea results support the idea that backbone hydrogen bonding plays a key role in difference in material states between IAssAI and AIssIA.

**Figure 4.**
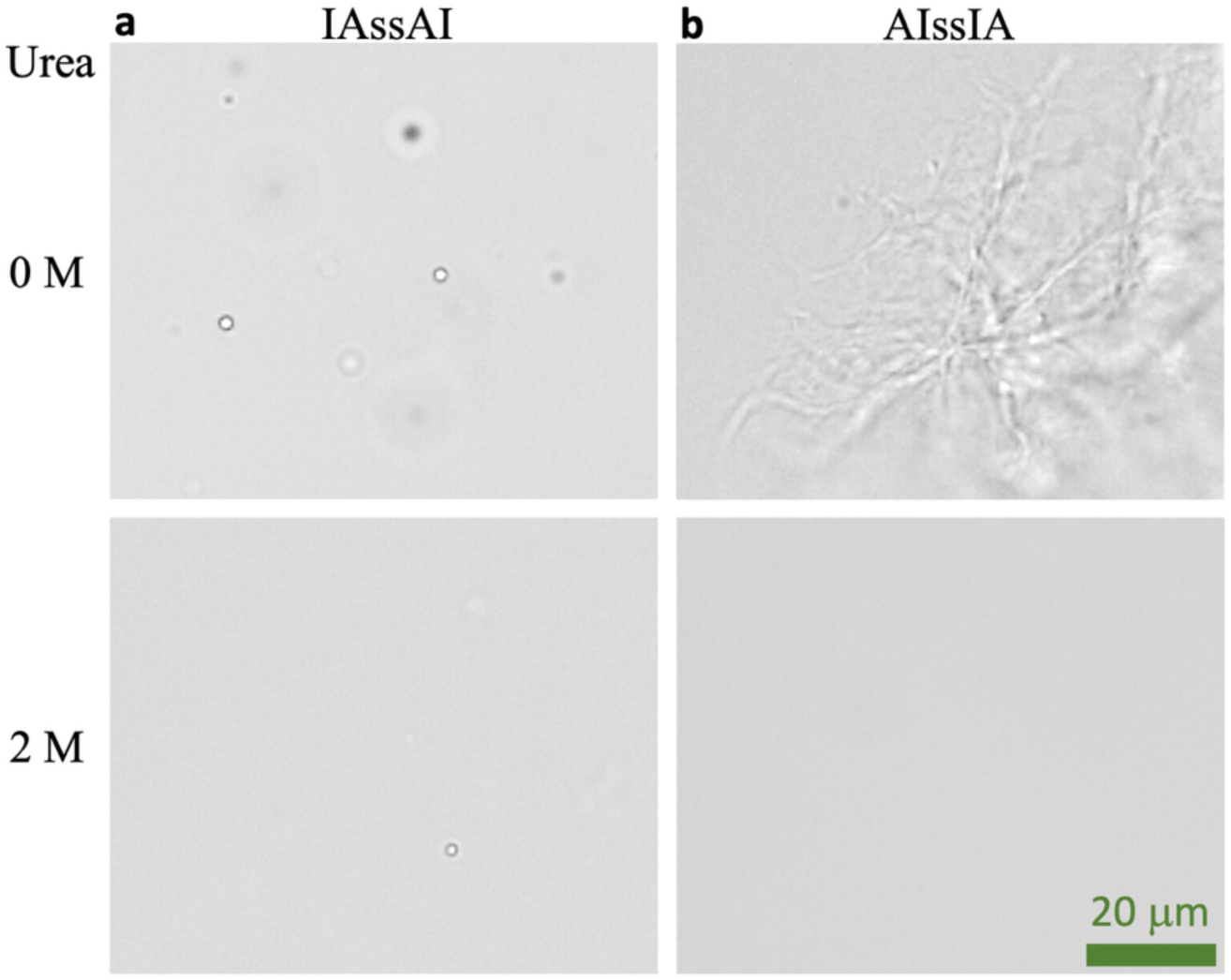
Urea dissolved AIssIA gels. (a, b) Brightfield images of 75 mg/mL IAssAI (panel a) and 7.5 mg/mL AIssIA (panel b), observed after preparation in Milli-Q water at pH 13 and 5-min incubation, without (top row) and with 2 M urea (bottom row).

### Addition of AAssAA converts AIssIA gels into droplets by reducing the level of hydrogen bonding

To further ascertain the role of backbone hydrogen bonding in gel formation, we sought to reduce the overall level of hydrogen bonding by mixing AIssIA with a low-hydrogen-bonding peptide, AAssAA. In MD simulations, an AIssIA:AAssAA mixture at a 1:1 molar ratio still forms gels, although, as expected, the average number of hydrogen bonds per half-chain is reduced from 0.46 to 0.37 (Figure 3c). When the molar ratio is changed to 1:4.3, the mixture now forms a flat slab (proxy for droplets), with the average number of hydrogen bonds per half-chain further reduced to 0.32. The MD simulations thus validated our design with AAssAA for diluting hydrogen bonding.

We also tested our design by determining the phase diagrams of AIssIA:AAssAA mixtures by brightfield microscopy. Pure AIssIA forms only gels (Figure 2e) whereas pure AAssAA forms amorphous dense liquids in most of the phase-separation region, except for a very small corner at very high pH (> 12) and high concentrations (≥ 200 mg/mL), where it forms gels ^23^. At a 1:2 molar ratio, AIssIA:AAssAA mixtures form amorphous dense liquids at pH up to 7 and concentrations up to 50 mg/mL, and form gels at higher pH and concentrations (Figure 5a). However, at a 1:5 molar ratio, while AIssIA:AAssAA mixtures still form amorphous dense liquids at low pH and concentrations, they form droplets at the corner with high pH and concentrations (Figure 5b). Therefore, diluting the hydrogen bonding of AIssIA with AAssAA produces a conversion from gels to droplets.

**Figure 5.**
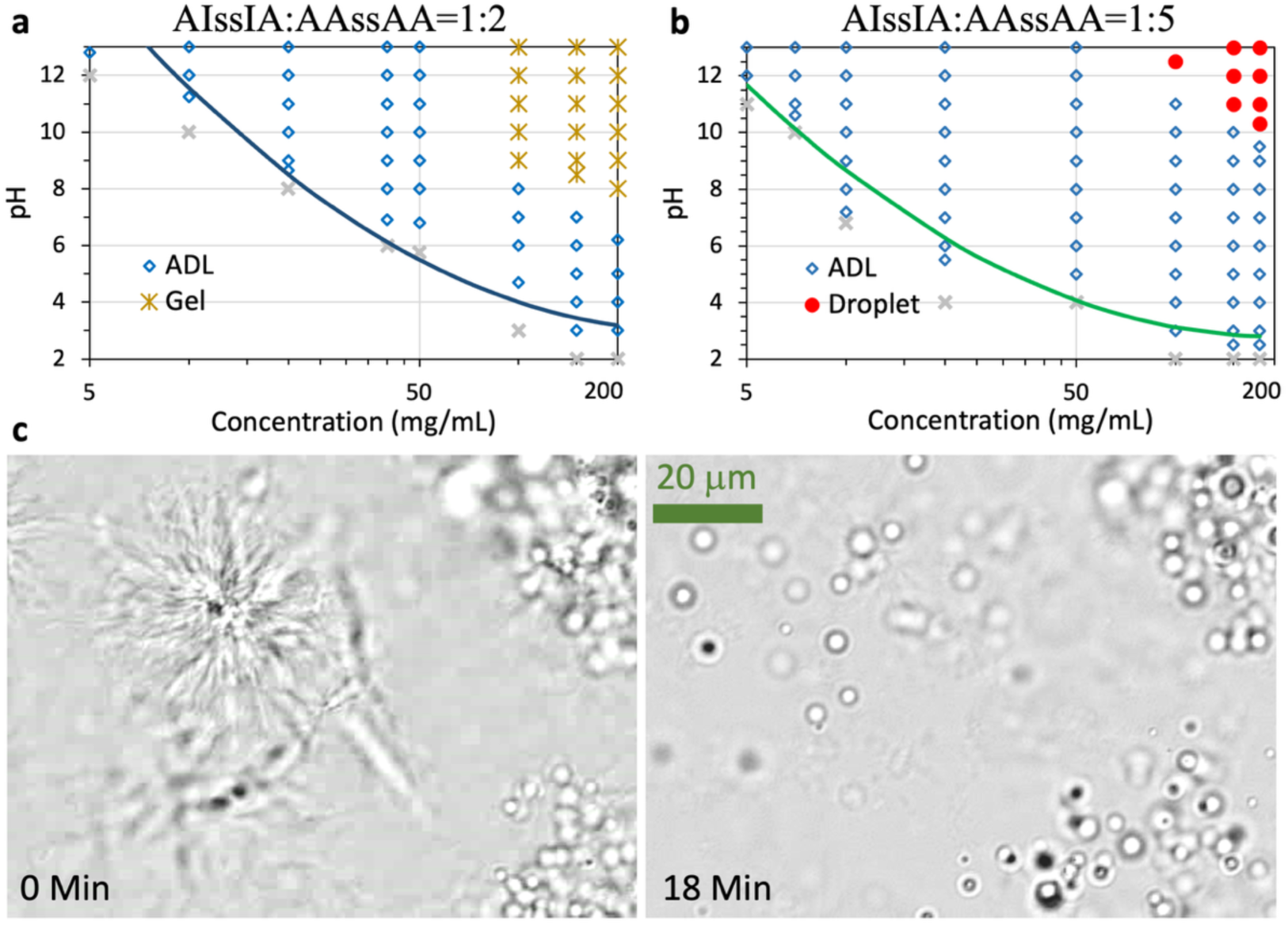
Addition of AAssAA converts AIssAI gels into droplets. (a, b) Phase diagrams of AIssIA:AAssAA mixtures at 1:2 (panel a) and 1:5 (panel b) molar ratios in 50 mM imidazole buffer. The abscissa displays the total concentration of the two peptides; phase diagrams were also confirmed after 2 hr of sample incubation. (c) Conversion of gels into droplets in a 2-μL aliquot of a 1:5 mixture at a 150 mg/mL total concentration in 50 mM imidazole (pH 13). The aliquot was loaded on a slide and observed continuously. A cropped region at 0 min (immediately after loading) and 18 min is shown. The entire 18-min video is presented as Video S2. The full field of view at 0 and 18 min is shown in Figure S1.

We directly captured this conversion by both brightfield and confocal microscopy. After raising the pH of an AIssIA:AAssAA sample (1:5 molar ratio; 150 mg/mL total concentration) to 13, a 2-μL aliquot was immediately placed on a glass slide and observed continuously under a brightfield microscope. At 0 min, droplets are observed in most of the field of view but gels also appear in some regions due to inhomogeneous mixing (Figure S1a; Figure 5c left panel). Over 18 min, as AAssAA floods in, gels shrink and droplets emerge (Video S2; Figure S1b and Figure 5c right panel). Similar conversion from gels to droplets is also observed on a confocal microscope (Figure S2; Video S3).

### N-methylation turns AIssIA gels into droplets or dense liquids by blocking backbone hydrogen bonding

To unequivocally establish that it is backbone hydrogen bonding that is promoting the gel formation of AIssIA, we synthesized N-methylated variants of AIssIA (see chemical structures in Figure S3 and Figure 6a, b). Methylation prevents an amide nitrogen from serving as a hydrogen bond donor. By selectively modifying either the inner or outer amide, we could potentially even pinpoint how hydrogen bonding at different sites in the peptide chain regulates material states. At 100 mg/mL and pH 13, outer N-methylated AIssIA (A_m_IssI_m_A) forms droplets but inner N-methylated AIssIA (AI_m_ss_m_IA) forms amorphous dense liquids (Figure 6a, b). Therefore, blocking hydrogen bonding at either the inner or outer nitrogen site disrupts gel formation. Still, the resulting condensates are different: droplets with outer methylation but amorphous dense liquids with inner methylation. This difference suggests a greater disruptive effect with inner methylation, consistent with the higher hydrogen bonding propensity of the inner site relative to the outer site seen in MD simulations (Figure 3c).

**Figure 6.**
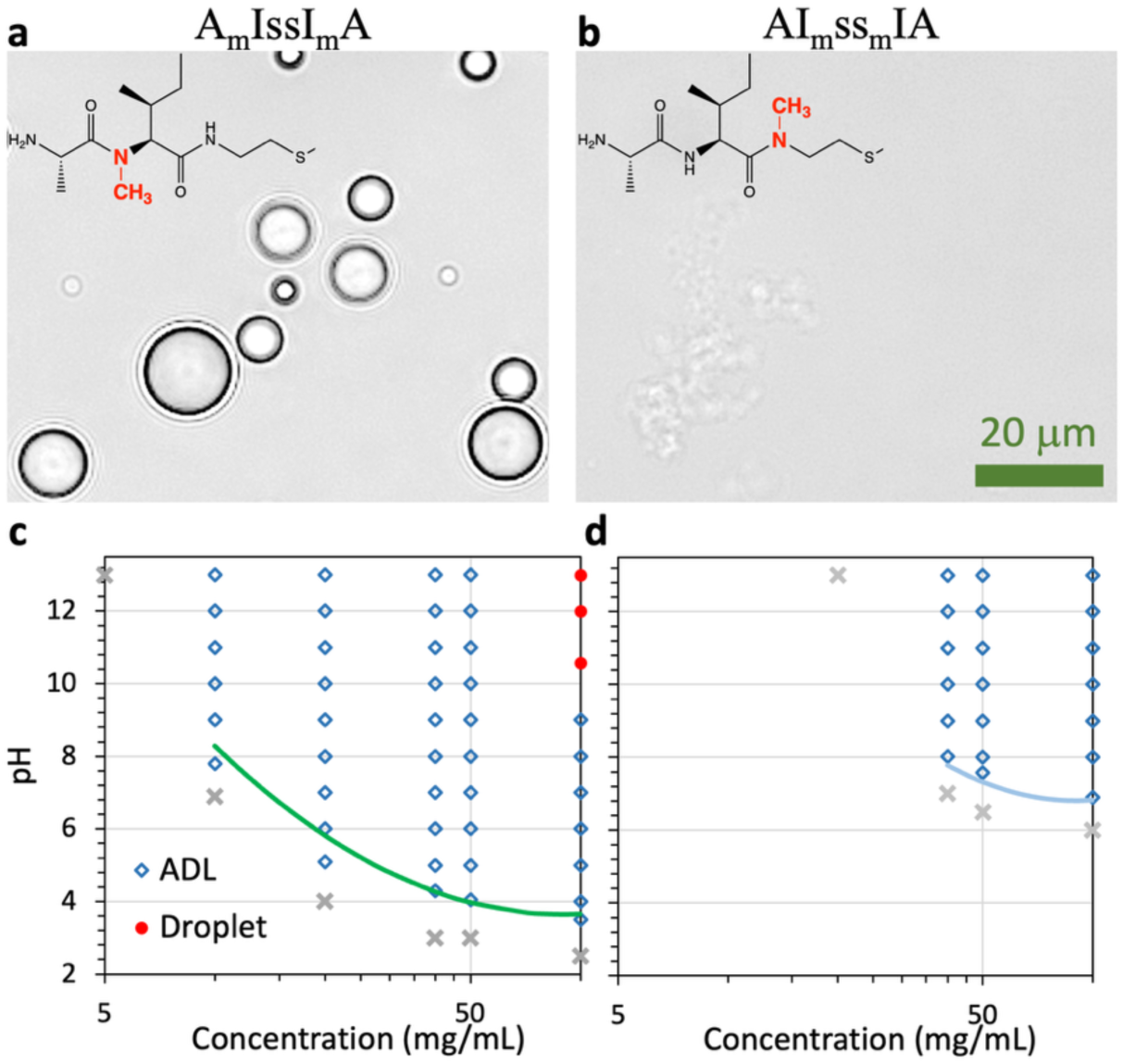
N-methylation of AIssIA turns gels into droplets or amorphous dense liquids. (a, b) Brightfield images of outer (panel a) and inner (panel b) N-methylated AIssIA samples at 100 mg/mL in Milli-Q water at pH 13. Superimposed on the brightfield images are chemical structures (for one half of the peptides) with methylation highlighted in red. (c, d) Phase diagrams of outer (panel c) and inner (panel d) N-methylated AIssIA in 50 mM imidazole buffer.

The phase diagram of A_m_IssI_m_A shows that it starts to phase separate at 10 mg/mL and forms amorphous dense liquids at concentrations up to 50 mg/mL (Figure 6c). Droplets form at 100 mg/mL and pH > 10. In comparison, AI_m_ss_m_IA has a higher threshold concentration, 40 mg/mL, and forms only amorphous dense liquids at concentrations up to 100 mg/mL (Figure 6d). The reduced tendency for AI_m_ss_m_IA to form droplets again indicates a greater disruptive effect with inner methylation. Overall, the phase diagrams demonstrate that hydrogen bonding at both the inner and outer sites is essential for the stability of gel networks, as methylation at either site changes the material state from gels to either droplets or amorphous dense liquids.

For comparison, we also methylated IAssAI. At 100 gm/mL and pH 13, I_m_AssA_m_I still forms droplets (Figure 7a, b) but IA_m_ss_m_AI only forms amorphous dense liquids (Figure 7c). However, droplets are formed when the IA_m_ss_m_AI concentration increases to 200 mg/mL (Figure 7d). These results suggest that backbone hydrogen bonding is not essential for forming droplets or amorphous dense liquids.

**Figure 7.**
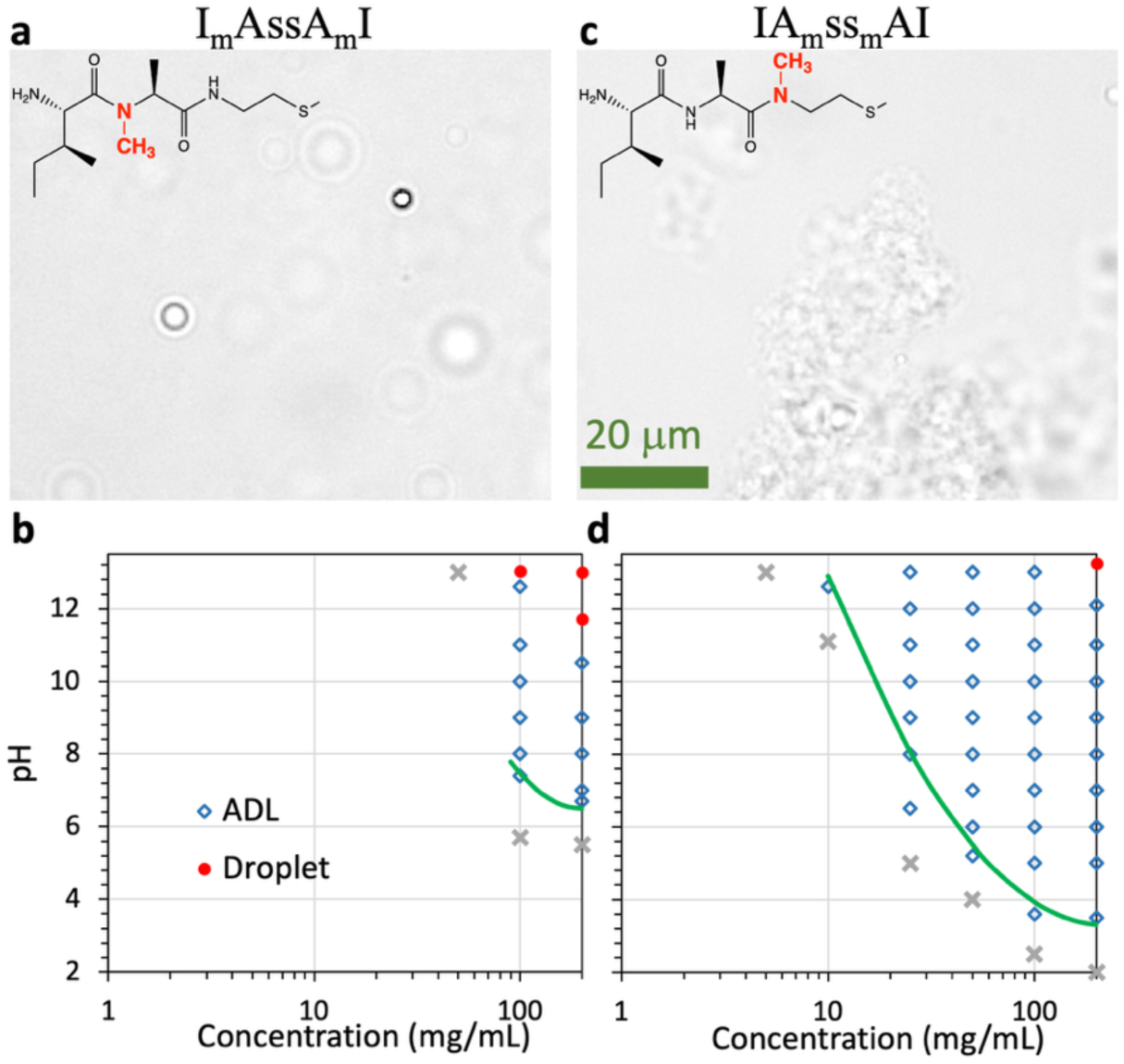
Material states of IAssAI remain the same after N-methylation. (a, b) Brightfield images of outer (panel a) and inner (panel b) N-methylated IAssAI at 100 mg/mL in Milli-Q water at pH 13. Superimposed on the brightfield images are chemical structures (for one half of the peptides) with methylation highlighted in red. (c, d) Phase diagrams of outer (panel c) and inner (panel d) N-methylated IAssAI in 50 mM imidazole buffer.

We simulated the spontaneous phase separation of the N-methylated AIssIA and IAssAI peptides. Under our simulation conditions, all four methylated peptides form a slab with rough surfaces, as illustrated by a snapshot of the I_m_AssA_m_I system in Figure 3d, leading us to classify them as amorphous dense liquids. As expected, the level of hydrogen bonding is reduced by N-methylation, to levels even lower than that of AAssAA (Figure 3c).

### NOESY reveals extensive interchain contacts and large molecular networks in AIssIA gels

To probe the structural and dynamic characteristics of intermolecular networks inside XYssYX condensates, we conducted ^1^H-^1^H NOESY experiments on a 500 MHz spectrometer. We looked for optimal conditions for the NMR experiments and settled on 10 and 15 mg/mL for AIssIA, 50 to 200 mg/mL for IAssAI, and 20 mg/mL for AAssAA, in Milli-Q water with 10% D_2_O at pH 13. Brightfield images confirmed that under these conditions, the three peptides form gels, droplets, and amorphous dense liquids, respectively (Figure S4). Given the lengthy time of the NMR experiments, one concern was condensate sedimentation (Figure S5). Gels, once formed, do not sediment due to network stability. Amorphous dense liquids sediment minimally, due to a very small difference in density from the bulk phase. Droplets, in contrast, exhibit a growing tendency of sedimentation as the peptide concentration increases, due to the increase in droplet size (see Figure S4b, c). We thus acquired ^1^H-^1^H NOESY spectra for 13 hrs on AIssIA gel samples, 50-mg/mL IAssAI droplet samples, and AAssAA dense-liquid samples, for 3 hrs on 100-mg/mL IAssAI droplet samples, and for 1 hr on 200-mg/mL IAssAI droplet samples. 1D NMR spectra of the three peptides, with peak assignments, are presented in Figure S6-S8.

We display the ^1^H-^1^H NOESY spectra of 10-mg AIssIA at pH 2 and pH 13 in Figure S9a, b and their zoomed views into one off-diagonal side in Figure 8a, b. ^1^H-^1^H NOEs can change signs depending on the effective tumbling time, which is approximately proportional to the molecule (or molecular cluster) that tumbles together with the two protons in question. ^1^H-^1^H NOEs have an opposite sign (referred to as negative hereafter) to the diagonal peaks for small molecular sizes but the same sign (or positive) as the diagonal peaks for larger molecular sizes. The crossover molecular weight (MW_x_) is approximately 900 Da for a 500 MHz spectrometer. At pH 2, AIssIA is a homogeneous solution and interchain interactions should essentially be absent. NOEs are detected between H_α_ and protons on side chains (Figure 8a; see Figure S3 for nomenclature). Most likely, these NOE contacts are formed within the same residue, where the proton pairs in question are separated by one to three covalent bonds between heavy atoms and thus have relatively short interatomic distances. Moreover, these NOEs have a negative sign, consistent with contacts within a single chain, which has a molecular weight of 520 Da that is below MW_x_.

**Figure 8.**
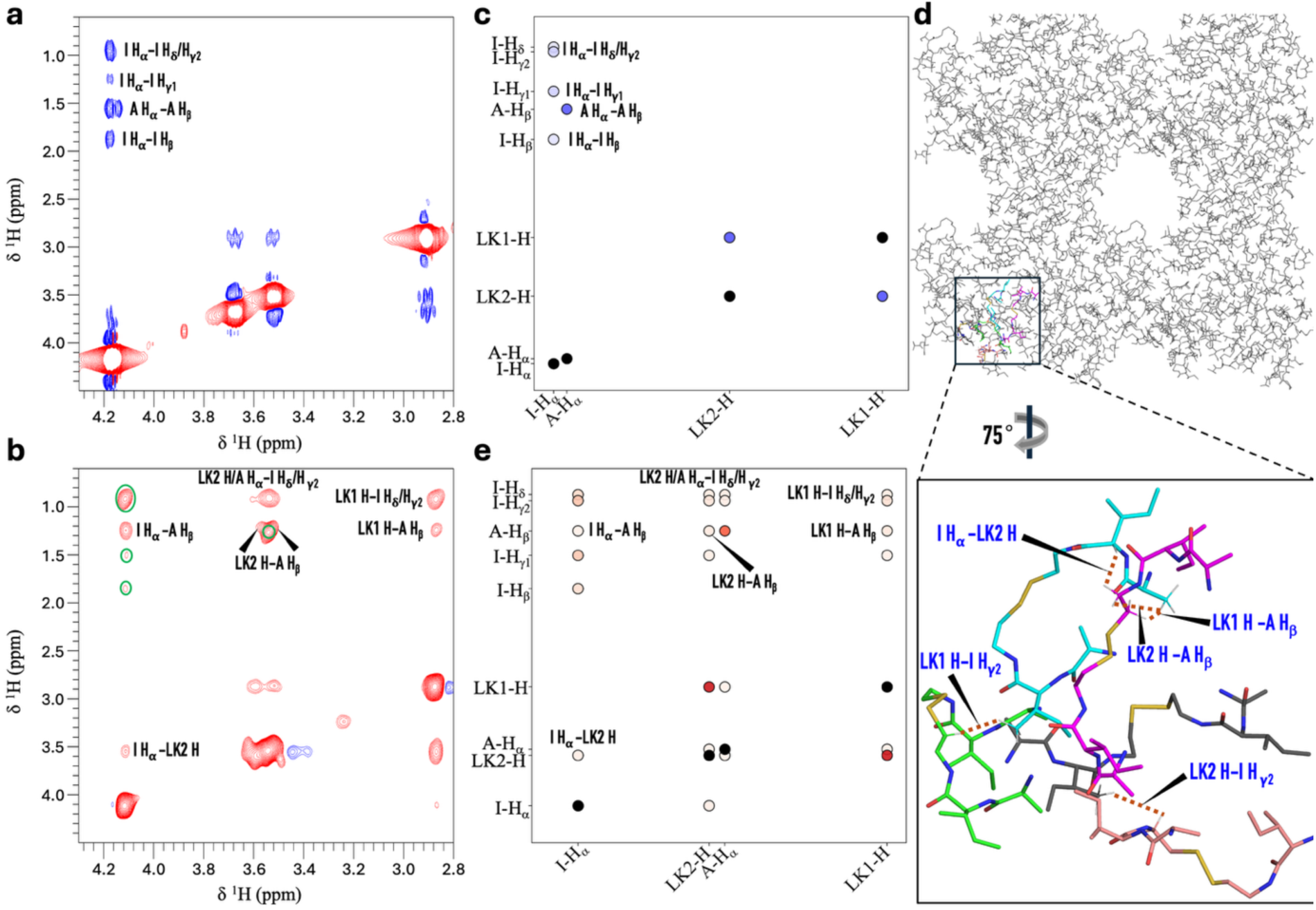
Intermolecular interactions in a gel-forming AIssIA sample. (a, b) ^1^H-^1^H NOESY spectra of 10 mg/mL AIssIA at pH 2 (panel a) and pH 13 (panel b) in Milli-Q water with 10% D_2_O. Spectra were acquired on a 500 MHz spectrometer with a 500-ms mixing time at 25 °C. NOEs with the same or opposite sign as the diagonal peaks are displayed as red and blue contours, respectively. (c) ^1^H-^1^H NOEs calculated from an MD simulation of AIssIA at a low-pH, dilute state. (d) Intermolecular interactions in AIssIA gel networks formed at high pH. A zoomed view highlights ^1^H-^1^H NOE contacts between AIssIA chains; two of these chains (with carbon in green and cyan) are also highlighted in Figure 3b. (e) ^1^H-^1^H NOEs calculated from an MD simulation of AIssIA at the high-pH, dense state illustrated in panel d. For (c, e), average *r*^-6^ values are displayed by the intensity of blue or red color; diagonal peaks are shown as black circles.

Upon gel formation at pH 13, many more NOEs are detected and they now all have a positive sign (Figure 8b). In addition to the NOEs that are already noted in the spectrum at pH 2 (indicated by green circles in Figure 8b, heavily contributed by proton pairs within the same residue), NOEs are detected for proton pairs that are separated by eight (LK1 H – A H_β_), seven (LK2 H – A H_β_), six or seven (LK1 H – I H_δ_/H_γ2_), five or six (LK2 H – I H_δ_/H_γ2_; A H_α_ – I H_δ_/H_γ2_) covalent bonds between heavy atoms. Given the corresponding long distances for these pairs within a chain, the latter NOEs likely have major contributions from interchain proton pairs. The positive sign of all the NOEs indicates that the NMR signals originate from chains that tumble together as a cluster, such that the total molecular weight exceeds MW_x_. The 10 mg/mL concentration is just inside the gel phase boundary, where only a small fraction of the peptide chains is in the gel phase while a majority of them are in the bulk phase. We speculate that chains in the bulk phase exchange with the gel phase, particularly with those at the surfaces of gel networks. After the exchange, a former bulk-phase chain becomes a part of a large molecular cluster, leading to a long apparent tumbling time (τ_app_) and hence positive NOEs.

We repeated the NOESY experiment using 15 mg/mL AIssIA. The results are qualitatively similar to those at 10 mg/mL, but fewer NOEs are detected upon gel formation (i.e., at pH 13; Figures S9c, d and S10a, b). The loss of some NOEs is due to line broadening. In Figure S10c, we overlay slices at 4.1 ppm (I H_α_ resonance) from the pH-13 NOESY spectra acquired from the 10-mg/mL and 15-mg/mL samples. The overlay shows line broadening of the I H_α_ resonance and loss of cross peaks including I H_α_ – I H_β_ at 15-mg/mL. Line broadening can be attributed to a further increase in τ_app_: at the higher concentration, gel networks become more connected, and each chain becomes a part of a larger molecular cluster, which could potentially grow into system-spanning. Indeed, all resonances are broadened beyond detection in a gel sample formed with 100 mg/mL AIssIA.

To validate the above interpretations of the NMR data, we simulated NOESY spectra by calculating average *r*^-6^ values in the MD trajectories, where *r* denotes the distance between proton pairs. We treat these average *r*^-6^ values as a proxy for NOEs. For the pH 2 condition, the trajectories used were those of a single AIssIA chain in solution, with both termini charged. The simulated NOESY spectrum (Figure 8c) indeed looks similar to the experimental counterpart (Figure 8a), therefore validating the assumption that NOEs at pH 2 are from proton pairs within a single chain. For the pH 13 condition, we calculated average *r*^-6^ values on the dense phase formed from 64 AIssIA chains with neutral termini (Figure 8d). The simulated NOESY spectrum at high pH (Figure 8e) again looks similar to the experimental counterpart (Figure 8b), therefore validating the assumption that NOEs at pH 13 are dominated by proton pairs within molecular networks in the gel phase. For most of the identified cross peaks in Figure 8e (same as in Figure 8b), > 75% of the average *r*^-6^ values are contributed by interchain proton pairs, as can be expected from the long distances of these pairs within a single chain (see above). Figure 8d presents a zoomed view of such interchain proton pairs. The simulated NOESY spectrum for a single AIssIA chain with neutral termini (modeling high pH; Figure S11) confirms that the cross peaks identified in Figure 8b cannot arise solely from intrachain proton pairs. Therefore, together with MD simulations, the NOESY data reveal both extensive interchain contacts and large molecular networks in AIssIA gels.

### Intermolecular association is more transient in IAssAI droplets

To compare and contrast with AIssIA gels, we also performed ^1^H-^1^H NOESY experiments on droplet-forming IAssAI samples. At 50 mg/mL and pH 13, IAssAI starts to form droplets (Figure 2d and Figure S4c). However, the ^1^H-^1^H NOESY spectra (Figure S12a-d) do not show the change in NOE sign as observed on the 10-mg AIssIA sample. Indeed, at both pH 2 and pH 13, only cross peaks from intrachain proton pairs are detected, such as between H_α_ and protons on a side chain (likely from the same residue). The intrachain attribution is confirmed by simulated NOESY spectra for a single IAssAI chain with charged or neutral termini (Figure S12e, f). The negative sign and intrachain attribution of the NOEs indicate that IAssAI tumbles effectively as individual chains in the 50-mg/mL sample at pH 13. Although each chain can spend part of its time inside droplets, those excursions are too transient to have an effect on NOEs.

When the IAssAI concentration is increased to 100 mg/mL, we see a switch in NOE sign from negative at pH 2 to positive at pH 13 (Figures S13a, b and S14a, b). Also similar to the 10-mg/mL AIssIA spectrum, many of the NOEs at pH 13, e.g., LK1 H – I H_δ_/H_γ2_, LK2 H – I H_δ_/H_γ2_, and LK1 H – I H_β_, are for proton pairs that have long distances within a chain and thus must arise from interchain contacts. Again, simulated NOESY spectra confirm the intrachain attribution at pH 2 (Figure S14c) and interchain attribution at pH 13 (Figure S14d). At the higher concentration, the total volume of droplets increases and hence a given chain now spends more time in the droplet phase, during which each chain is a part of a large molecular cluster. Those times make a dominant contribution to the observed NOEs, explaining their interchain origin and positive sign. The NOEs of the 200-mg/mL IAssAI sample also show a sign switch from negative at pH 2 to positive at pH 13, although sedimentation makes it difficult to observe all the cross peaks (Figure S13c, d).

The concentrations required for a chain to be detected by ^1^H-^1^H NOESY as a part of a large molecular cluster are very different for gel-forming and droplet-forming samples: 10 mg/mL for AIssIA and 100 mg/mL for IAssAI. We infer that AIssIA gel networks comprise strong, backbone hydrogen bonding-buttressed molecular clusters. In comparison, backbone hydrogen bonding plays a lesser role, and hydrophobic interaction-dominated intermolecular association is more transient in IAssAI droplets. Amorphous dense liquids are characterized by very weak interactions. Consequently, the ^1^H-^1^H NOESY spectra of 20-mg/mL AAssAA show the same negative sign of NOEs upon a pH change from 2 to 13 and only intra-residue cross peaks at both pH values (Figure S15).

### Backbone hydrogen bonding likely drives directional growth of molecular networks

As illustrated by the MD snapshot in Figure 3b, a typical interchain hydrogen-bonding configuration involves two half-chains in parallel or antiparallel alignment, similar to a two-strand β-sheet but with the hydrogen bonds not necessarily in an alternating pattern. This analogy made us realize that the high hydrogen-bonding propensities of IIssII and VVssVV (Figure 3c) originate from the fact that the β-branched amino acids I and V are preferred in β-sheets, due to the ability of the β-branches in maintaining the rigidity of β-sheets. The preference is modest at the first position (on the N-terminal side) of β-strands and becomes strong starting at the second position ^41^. This positional dependence explains why AIsssIA has a higher hydrogen-bonding propensity than IAssAI.

We further observed that when two half-chains form hydrogen bonds, the two other half-chains are usually directed away (Figure 3b), to avoid clashes and also to interact with additional chains. As a result, chains forming hydrogen bonds are relatively straight. To quantify this tendency, we calculated the angle between the two halves of each chain (Figure 9a). The probability distribution of the intrachain angle for AIssIA in the dense phase is shown in Figure 9b. The peptide forms 0.51 hydrogen bonds per half-chain when the intrachain angle is > 120° but only 0.38 hydrogen bonds per half-chain when the intrachain angle is < 120° (Figure 9c). We measured the chain straightness by the probability ratio of the intrachain angle being > 120° over being < 120° (Figure 9b). This probability ratio ranges from 0.71 to 0.92 in the gel phase of VVssVV, IIssII, and AIssIA, but spans a lower range, from 0.51 to 0.60, in the droplet phase of IAssAI and LLssLL (Figure 9d). In short, interchain hydrogen bonding requires peptide chains to be relatively straight. Hydrophobic interactions, in contrast, do not have this requirement.

**Figure 9.**
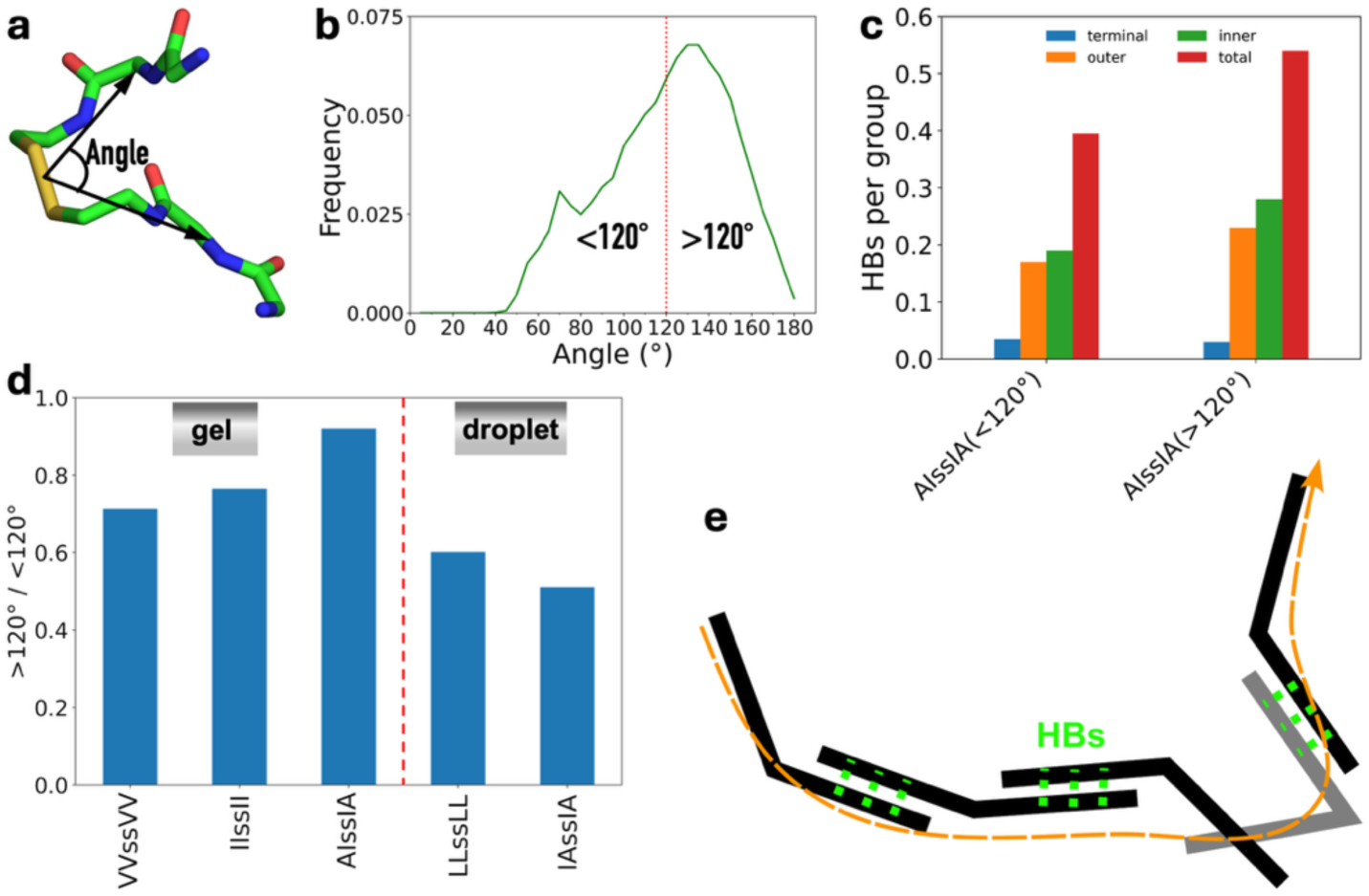
Interchain hydrogen bonding requires peptide chains to be relatively straight. (a) The angle between the two halves of a peptide chain. (b) Probability distribution of the intrachain angle for AIssIA in the dense phase. (c) Average number of hydrogen bonds per half-chain in the AIssIA dense phase, broken down according to bent (< 120°) or straight (> 120°) angles. (d) Probability ratio of straight and bent chains in the dense phase. (e) Molecular network growth via backbone hydrogen bonds (HBs; black chains) or hydrophobic interactions (gray chain). An orange curve indicates growth direction.

When chains are relatively straight, molecular networks will grow in a fixed direction (Figure 9e). The direction can change when a bent chain is added via forming hydrophobic interactions. Directional growth can be a defining feature of gels (Figure 1c).

## Discussion

By combining optical microscopy, NMR spectroscopy, and all-atom MD simulations, we have thoroughly investigated backbone hydrogen bonding as a determinant of material states for XYssYX condensates, and established that it promotes gel formation of AIssIA. We have taken three routes to reduce hydrogen bonding levels: adding urea, mixing with a low-hydrogen-bonding peptide, and N-methylation. All three routes result in the dissolution of gels or conversion to droplets or amorphous dense liquids. ^1^H-^1^H NOESY and MD simulations together have revealed that backbone hydrogen bonding reinforces molecular networks and likely drives their directional growth.

The presented study was motivated by the puzzling observation that LLssLL and MMssMM formed droplets but IIssII and VVssVV formed gels ^23^. Adding to the intrigue, we initially found that IAssAI formed droplets but AIssIA formed gels. MD simulations showed high hydrogen-bonding levels in all the gel-forming peptides (Figure 3c). By inspecting the hydrogen-bonding configurations in AIssIA gels (Figure 3b), we noticed that they bear resemblance to β-sheets. We finally realized that the gel-forming peptides all have I or V at the inner positions; at inner positions, amino acids with β-branches, including I and V, stabilize β-sheets by maintaining their rigidity. Therefore, the ability of I/V-containing peptides is ultimately traced to β-branching. Of course, the side chains of these amino acids also form hydrophobic interactions. Conversely, droplet-forming peptides can also form backbone hydrogen bonds. All favorable interactions contribute to condensate formation ^7^, but we now see that the types of interactions can tune their material states.

Another important observation from MD simulations is that interchain hydrogen bonding leads to relatively straight peptide chains (Figure 9c). Networks of straight chains tend to grow directionally (e.g., Figure 9e). We recognize that droplets, reversible aggregates, and amorphous dense liquids all grow isotropically, i.e., without a preferred direction (Figure 1a, b, d); in contrast, gels are distinct in being filamentous (Figures 1c and 4b), suggesting directional growth. For I/V-containing peptides, backbone hydrogen bonding acts as the link between β-branching and directional growth. Gels are the least understood material state; the insights gained here will narrow some of this gap and be instructive for unraveling the mysteries behind other gels. For example, Abbas et al. ^34^ observed that YFsFY and YFoFY formed droplets but YFcFY formed gels. One possible explanation is that the -C- linker maintains straighter chain conformations relative to the -S- and -O- linkers.

Our integrative approach presents significant opportunities for in-depth investigation of customizable peptides, where subtle chemical changes can be used to tune condensate material states. Substitution of I with L, change in the order between A and I along the sequence, and N-methylation are examples where the material state is changed from gels to droplets. Manipulating backbone hydrogen bonds in addition to side-chain interactions will enrich the toolbox for condensate engineering.

### Experimental Section

#### Materials

Boc-N-methyl-L-alanine (catalog # 15549), Boc-N-methyl-L-isoleucine (catalog # CH6H9A56B61F), Boc-Ala-OH (catalog # 15380), Boc-Ile-OH (catalog # 359653), 1-hydroxybenzotriazole hydrate (HOBt; catalog # 157260), HBTU (catalog # 12804), N,N-diisopropylethylamine (DIPEA; catalog # 387649), cystamine dihydrochloride (CDC; catalog # 30050), and hydrogen chloride solution (4 M in dioxane; catalog # 345547) were from Millipore Sigma. 2,2’-Disulfanediylbis(N-methylethan-1-amine) dihydrochloride (catalog # AR01RJIL) was from Aaron Chemicals. All solvents were from Sigma Aldrich.

#### Synthesis

Synthesis of XYssYX peptides followed a similar procedure to one reported for XXssXX ^23^ and consisted of the following major steps. (1) *Amide coupling* to produce Boc-YssY-Boc (Y = A or I). Boc-Y-OH (1.55 mmol, 1 equiv), HOBt (1.47 mmol, 0.95 equiv), and HBTU (1.47 mmol, 0.95 equiv) were mixed in DMF (3 mL) in a round-bottom flask. DIPEA (6.2 mmol, 4 equiv, in 1.05 mL) and CDC (0.7 mmol, 0.45 equiv) were then added to the mixture. The reaction proceeded at room temperature for 24 hr and was monitored by thin-layer chromatography until completion. After the reaction, 40 mL of deionized (DI) water was added to the flask. The resulting precipitate of Boc-IssI-Boc was collected by centrifugation (4500 RPM for 30 min) in a 50 mL tube. Boc-AssA-Boc did not precipitate ad thus was separated using column chromatography with a CH₂Cl₂/MeOH mobile phase in a 15:1 ratio and then using a rotary evaporator to obtain Boc-AssA-Boc powder. (2) *Hydrolysis* to yield YssY. The Boc-YssY-Boc product was dissolved in dioxane/dichloromethane (DCM) (4:1 ratio, 1.5 mL) in a round-bottom flask. Hydrogen chloride (4 M in dioxane, 3 mL) was added, and the reaction was maintained at room temperature for 3 hr. The mixture was then concentrated using a rotary evaporator, and 40 mL of diethyl ether was added to induce precipitation. The white precipitate (YssY) was collected by centrifugation. (3) *Second round of amide coupling and hydrolysis* to synthesize XYssYX from YssY. Starting with YssY, amide coupling using Boc-X-OH yielded Boc-XYssYX-Boc. Hydrolysis yielded the final compound, XYssYX.

Synthesis of N-methylated peptides followed the same procedure with minor modifications. For outer N-methylation, Boc-N-methyl-L-alanine and Boc-N-methyl-L-isoleucine were used. For inner N-methylation, 2,2’-disulfanediylbis(N-methylethan-1-amine) dihydrochloride was used. The amide coupling reaction was conducted at 60 °C for 24 hr.

#### Purification and characterization

Final peptides were purified using a preparative HPLC system (Waters Prep 2545) equipped with a C18 column (Waters XBridge Peptide BEH). Following purification, samples were lyophilized to remove water, yielding a white powder. The chemical structure and purity of each peptide were confirmed by 1D ¹H NMR spectroscopy.

#### Preparation of peptide condensate samples

A stock solution was prepared by dissolving purified peptide powder in Milli-Q water or 50 mM imidazole buffer at room temperature. 37 % (w/w) HCl or 5 M NaOH was added to change pH. All solutions and buffers were filtered (0.22-μm pore size; Millipore Sigma catalog # SLGPR33RS) before use.

#### Brightfield microscopy

2-μL aliquots of peptide samples were placed on a glass slide and imaged under an Olympus BX53 microscope using a 40× objective at room temperature. Unless otherwise indicated, samples were imaged immediately after changing to the desired pH. Displayed images were processed using ImageJ.

#### Confocal microscopy

5-μL aliquots of peptide samples were placed into a custom sample holder and scanned using the confocal module of a LUMICKS C-Trap instrument at room temperature. The samples were doped with a viscosity-sensitive dye ^23, 42^ at 2 μM. A 20.6 μm ′ 27.6 μm field of view was scanned with excitation wavelength at 532 nm (10% laser power; 200 nm pixel size; 0.05 ms pixel dwell time).

#### NMR spectroscopy

Peptides were dissolved in Milli-Q water with 10% D_2_O (Cambridge Isotrope Laboratories catalog # DLM-4-100) at a total volume of 600 μL in 1.5-mL microcentrifuge tubes. The initial samples were at pH close to 2 and directly transferred to 5-mm Wilmad® NMR tubes (Millipore Sigma catalog Z272019) for ^1^H-^1^H NOESY. After acquisition for pH 2, the pH of the samples was raised to 13 by adding a known amount of 5-M NaOH (in 10% D_2_O) for acquisition for pH 13.

All NMR spectra were collected on a Bruker Advance III HD 500 MHz spectrometer at 25 °C. 1D NMR spectra were acquired using 32 scans with water suppression. For ^1^H-^1^H NOESY, the mixing time was 500 ms. The total acquisition times were: 10 and 15 mg/mL AIssIA, 50 mg/mL IAssAI, and 20 mg/mL AAssAA, 13 hr (with water suppression); 100 mg/mL IAssAI, 3 hr; 200 mg/mL IAssAI, 1 hr. For NMR data analysis, MestReNova was used for 1D NMR; NMRFX for ^1^H-^1^H NOESY, and Topspin 3.6.5 NMR was used for 1D spectral slices.

#### Molecular dynamics simulations of peptide phase separation

We conducted simulations of XYssYX, where X and Y were I or A. The default force field for the peptides was Amberff14SB ^43^; parameters for the half-linker (-NH-CH_2_-CH_2_-S-) were generated previously ^23^ using Amber tools ^44^ and Gaussian 16 ^45^. We also studied inner and outer N-methylated peptides. Specifically, we replaced the N-H groups with N-CH_3_ in inner (XY_m_ss_m_YX) or outer (X_m_YssY_m_X) positions using VMD ^46^ and patched the resulting residue with methylamine capping. The water model was TIP4PD ^47^.

Phase-separation simulations were carried out for six pure peptides (AIssIA, IAssAI, A_m_IssI_m_A, AI_m_ss_m_IA, I_m_AssA_m_I, and IA_m_ss_m_AI) as well as two AIssIA:AAssAA mixtures (at ratios 1:1 and 1:4.3). The simulations started with a random distribution of 64 chains, all with neutral termini (modelling high pH). For AIssIA and IAssIA, an 8-chain system was built first. The 8 chains were solvated with 623-627 waters in a cubic box of 30 Å side length. After energy minimization (2,000 steps of steepest descent and 3,000 steps of conjugate gradient, using sander), the system was simulated for 100 ps with a 1-fs timestep at constant NVT on GPUs using pmemd.cuda ^48^. The temperature was ramped from 0 to 294 K over the first 40 ps and maintained at 294 K for the remaining 60 ps. The last snapshot was duplicated in each of the three orthogonal directions to build a 64-chain system. The A_m_IssI_m_A and AI_m_ss_m_IA systems were prepared from the AIssIA system, by replacing AIssIA chains with the methylated counterparts. I_m_AssA_m_I and IA_m_ss_m_AI were prepared in a similar way from the IAssAI system. The AIssIA:AAssAA mixture at 1:1 molar ratio was prepared by randomly placing 32 AIssIA chains and 32 AAssAA chains in a cubic box with a side length of 60 Å. Similarly the 1:4.3 mixture was prepared using 12 AIssIA chains and 52 AAssAA chains. In all cases, the 64 peptide chains were solvated with 4600-5200 water molecules.

To prepare each system for condensation, 1/4, 1/3, 1/2, or 2/3 water molecules were removed randomly from the 60 Å cubic box, yielding four peptide concentrations ^23, 40^. After energy minimization for 5,000 steps, 500-ps equilibration was performed with a 1-fs timestep at constant NVT, where the temperature was ramped from 0 to 294 K for 40 ps and then maintained at 294 K for 460 ps. The system at each concentration was then simulated in duplicates at constant NPT (pressure at 1 bar and temperature at 294 K) for 4-6.5 ms with a 2-fs timestep.

For simulating the dilute phase, a single peptide chain was solvated with ∼6800 water molecules in a cubic box with a side length of 60 Å. Both charged (modelling low pH) and neutral (modelling high pH) forms of AIssIA, IAssAI, and AAssAA were simulated in the dilute phase. Each system was energy minimized for 5000 steps and equilibrated for 500 ps with at a 1-fs timestep at constant NVT, with temperature ramping from 0 to 294 K for 40 ps and maintained for 460 ps at 294 K. The simulations then continued at constant NPT for 1 ms with a 2-fs timestep.

All the simulations were run in AMBER22 ^49^. The particle mesh Ewald (PME) method ^50^ was employed to handle long-range electrostatic interactions using a nonbonded cutoff distance of 10 Å. The Langevin thermostat with a damping constant of 3 ps^-1^ was used to regulate temperature, while the Berendsen barostat ^51^ was used for pressure regulation. All bonds involving hydrogen atoms were constrained using the SHAKE algorithm ^52^.

#### MD data analyses

Interchain hydrogen bonds and intrachain angles were calculated using CPPTRAJ ^53^ over a 1,000-ns portion of phase-separation simulations sampled at 1-ns intervals. In the chosen 1000-ns portions, the peptides formed flat slabs (proxy for droplets), slabs with holes (proxy for gels), and slabs with rough surfaces (proxy for amorphous dense liquids). Hydrogen bonds were defined with a 3.0-Å distance cutoff and a 135° angle threshold; hydrogen bond calculations were performed after centering the periodic system on each peptide chain. The intrachain angle was defined between two vectors that originated from the center of the two sulfur atoms, each ending at the center of two carbonyl carbon atoms in a half-chain. Hydrogen bond and intrachain angle results were averaged over the 64 chains.

For NOE calculations, MDAnalysis ^54^ and in-house python scripts were used. For the dense phase, calculations were done on the foregoing 1,000-ns portions but sampled at 100-ns intervals. For the dilute phase, NOE calculations were also done on 1000-ns portions of single-chain simulations. NOEs were normalized by the number of peptide chains. VMD and Pymol (https://pymol.org) were utilized for visualization and rendering images.

## Acknowledgments

We thank Dr. Daniel J. McElheny for assistance with NMR experiments, Dr. McElheny and Dr. Justin L. Lorieau for discussion, Dr. Daesung Lee for assistance with peptide synthesis, and Dr. Andy I. Nguyen for giving us D_2_O. This work was supported by National Institutes of Health Grant GM118091.

---

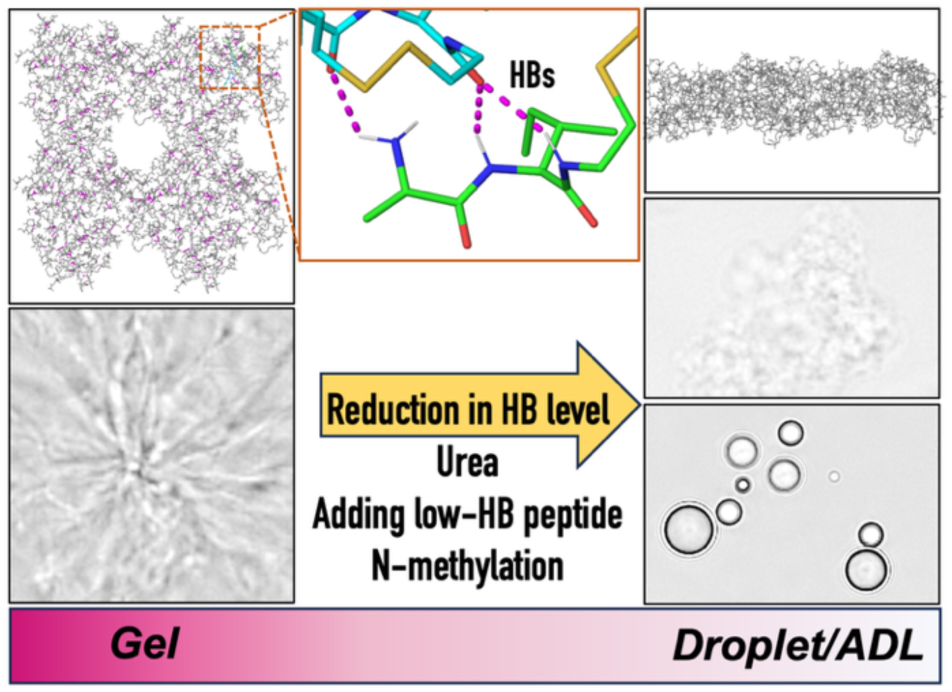

