## Supporting Figures for "Backbone Hydrogen Bonding as a Determinant of Condensate Material States"

*Supporting information*

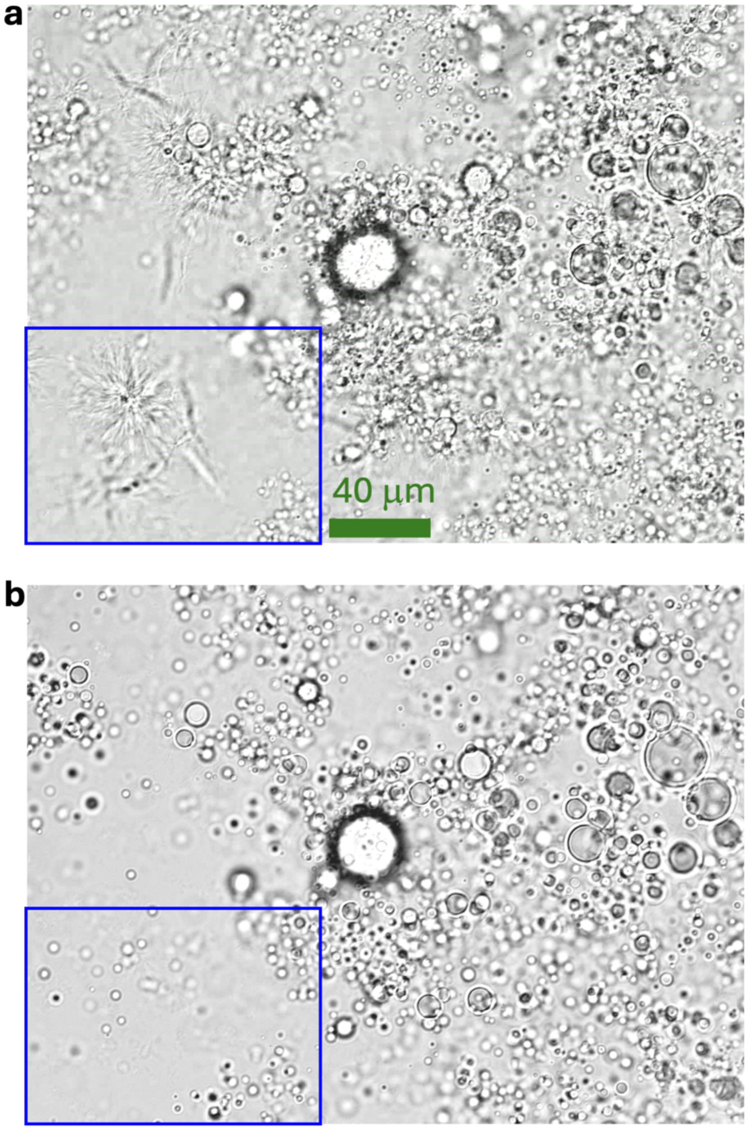


Figure S1. Full view of Figure 5c. Brightfield images of an aliquot of an AIssIA:AAssAA mixture placed on a slide, observed at (a) 0 and (b) 18 min. The mixture was at a 1:5 molar ratio and a 150 mg/mL total concentration in 50 mM imidazole (pH 13).

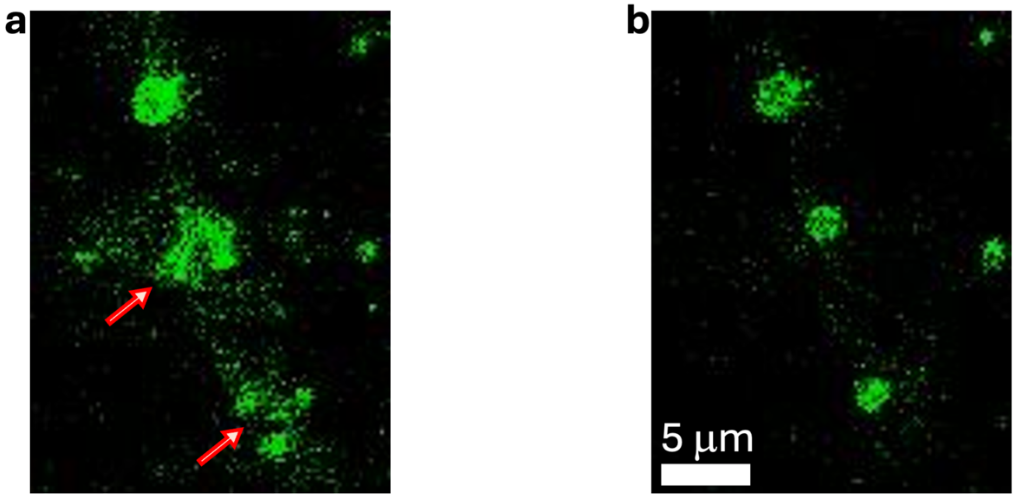


Figure S2. Confocal images of an AIssIA:AAssAA mixture (molar ratio 1:5) at a 150 mg/mL total concentration in 50 mM imidazole (pH 13). The sample was stained with a viscosity-sensitive dye (2 µM). A 5-µL aliquot was pipetted into a custom sample holder and imaged on the confocal module of a LUMICKS C-Trap instrument. In (a), arrows point to gels, which are converted to droplets in (b).

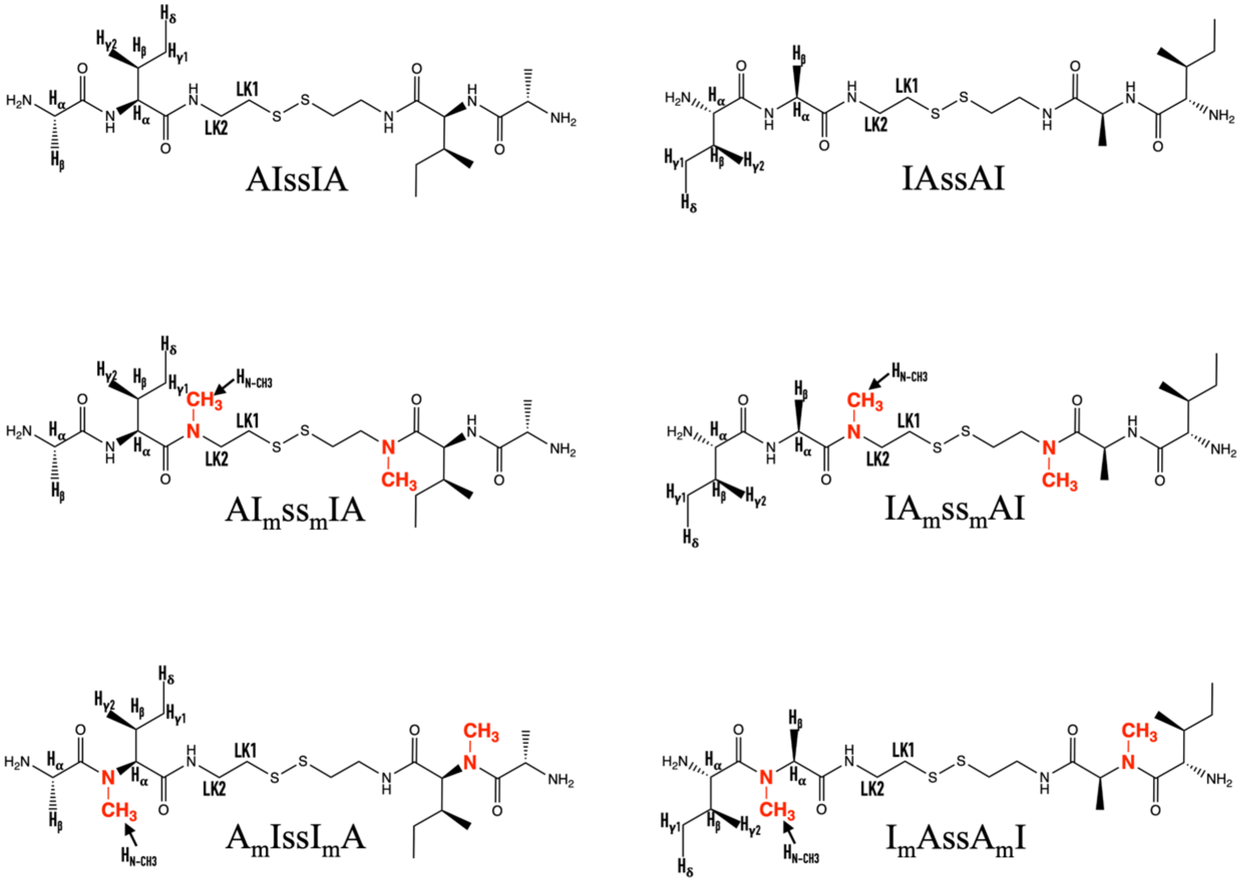


Figure S3. Chemical structure of AIssIA and IAssAI and their N-methylated variants.

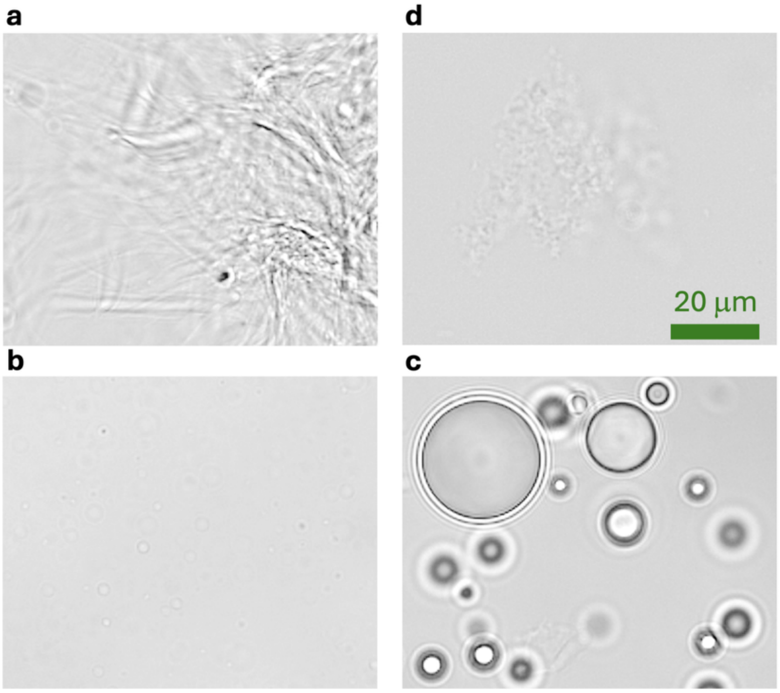


Figure S4. Brightfield images of (a) 10 mg/mL AIssIA, (b) 50 mg/mL IAssAI, (c) 100 mg/mL IAssAI, and (d) 20 mg/mL AAssAA. Samples were prepared in Milli-Q water with 10% $D_2O$ at pH 13.

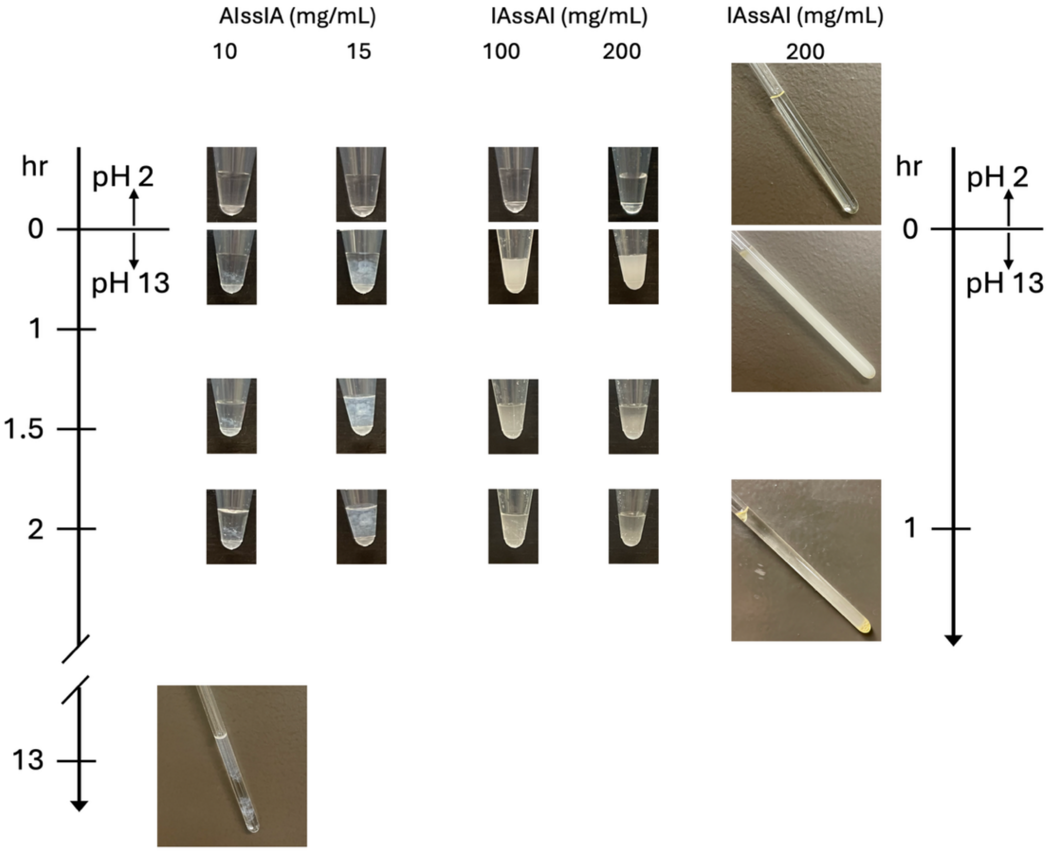


Figure S5. Images of 30-µL samples of AIssIA and IAssAI in microtubes and 500-µL samples in 5-mm NMR tubes. Images were taken when the samples were initially at pH 2 and at various time points after raising pH to 13. Samples were prepared in Milli-Q water with 10% $D_2O$ at pH 13.

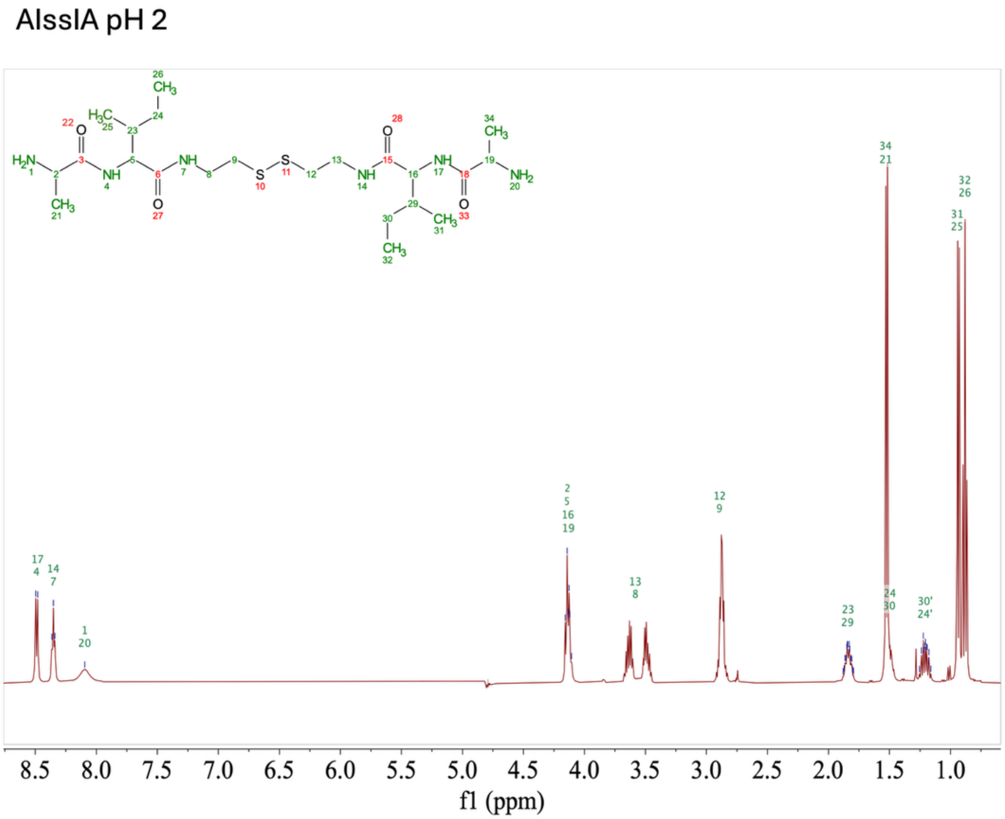


Figure S6. [1]H NMR spectrum of 10 mg/mL AIssIA in Milli-Q water with10% $D_2O$ at pH 2. Primed and unprimed labels for the same site indicate peak splitting.

IAssAI pH 2

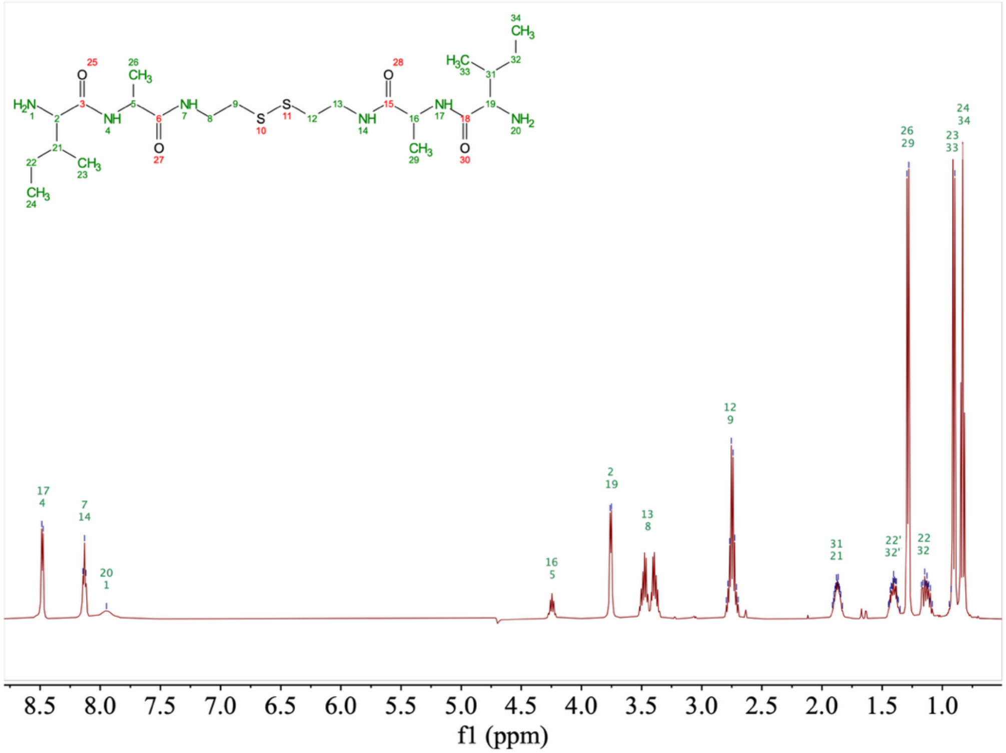

Figure S7. [1]H NMR spectrum of 100 mg/mL IAssAI in Milli-Q water with10% $D_2O$ at pH 2. Primed and unprimed labels for the same site indicate peak splitting.

AAssAA pH 2

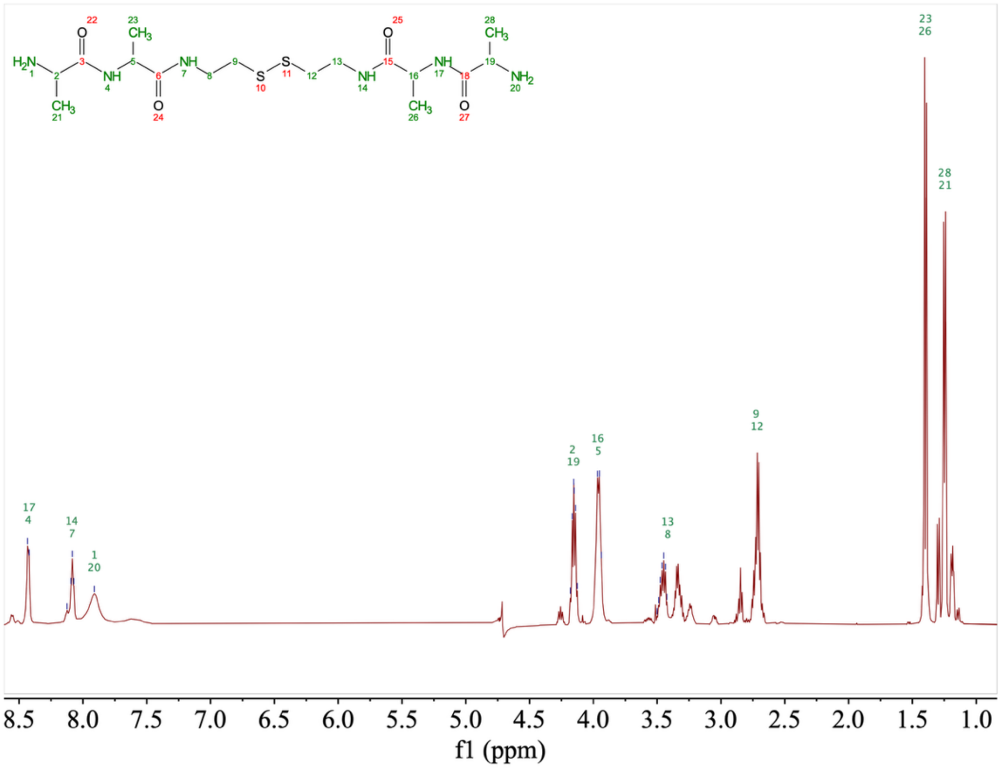


Figure S8. $^{1}H$ NMR spectrum of 20 mg/mL AAssAA in Milli-Q water with10% $D_2O$ at pH 2.

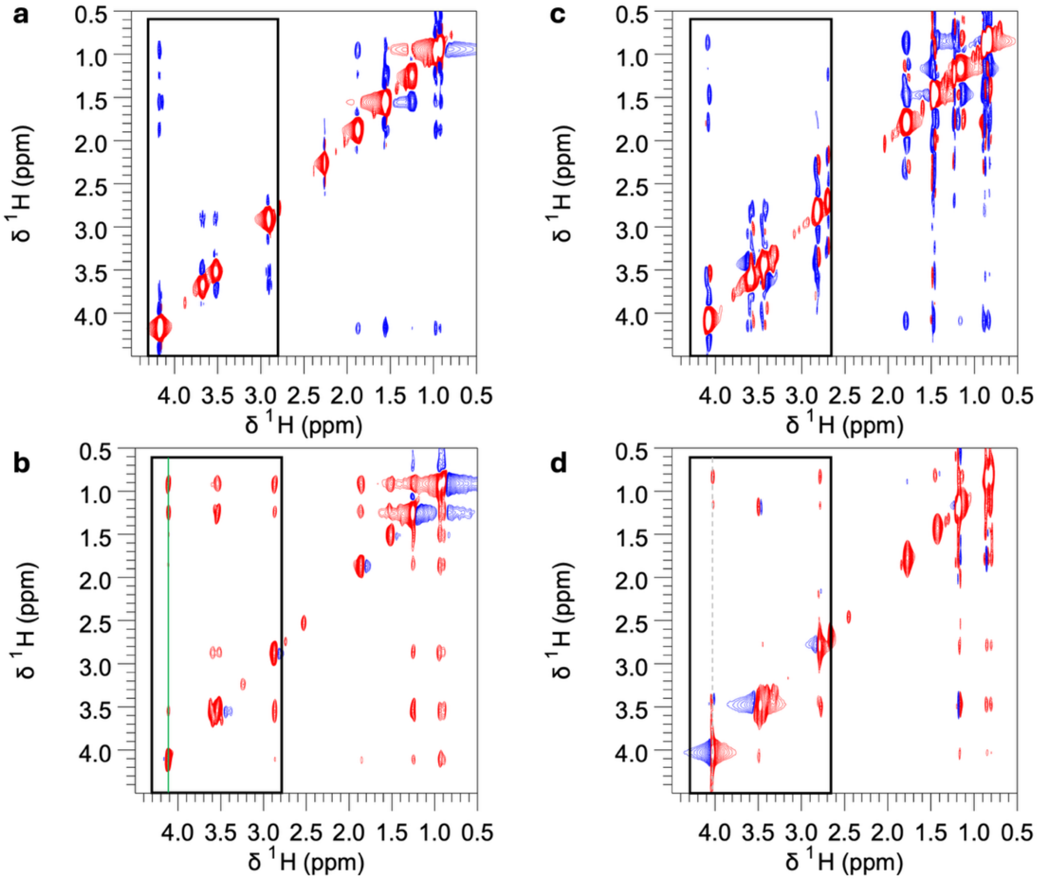


Figure S9. $^1H$-$^1H$ NOESY spectra of AIssIA, at 10 mg/mL and (a) pH 2 and (b) 13, or at 15 mg/mL and (c) pH 2 and (d) 13. Samples were prepared in Milli-Q water with 10% $D_2O$; spectra were acquired on a 500 MHz spectrometer with a 500-ms mixing time at 25 °C. Black rectangular boxes in (b) and (d) indicate regions that are shown in an enlarged view in Figure 8a, b and Figure S10a, b, respectively. A green solid line in (b) and a gray dashed line in (d) are drawn at the I $H_\alpha$ frequency to mark the 1D slices shown in Figure S10c. In (b) and (d), blue contours are artifacts arising from imperfect phasing.

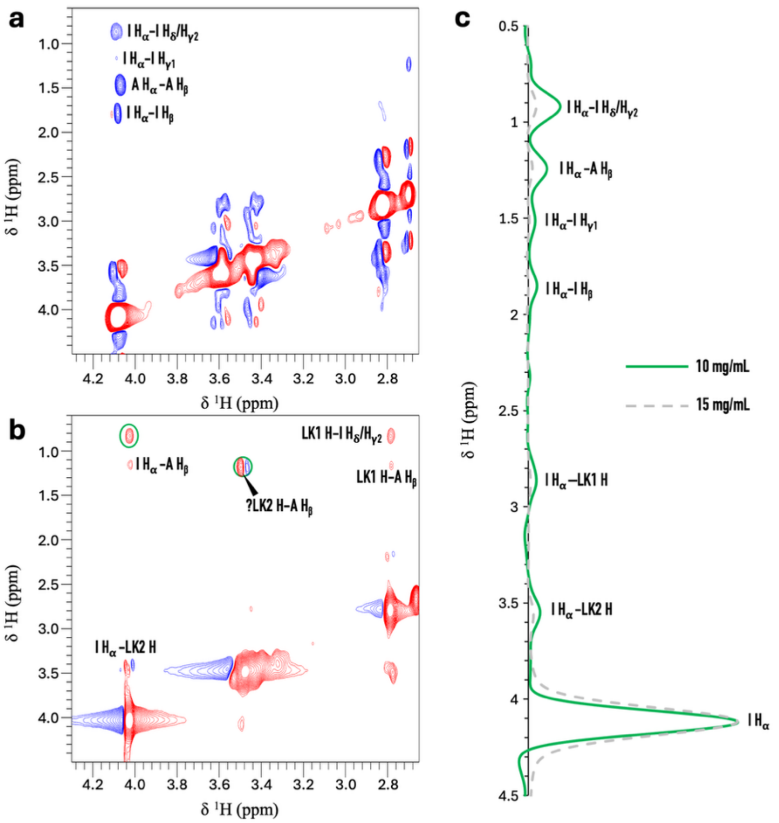


Figure S10. Enlarged view and 1D slices of Figure S9. (a, b) Enlarged view of the boxed regions in Figure S9c, d. (c) 1D slices through the lines marked in Figure S9b, d.

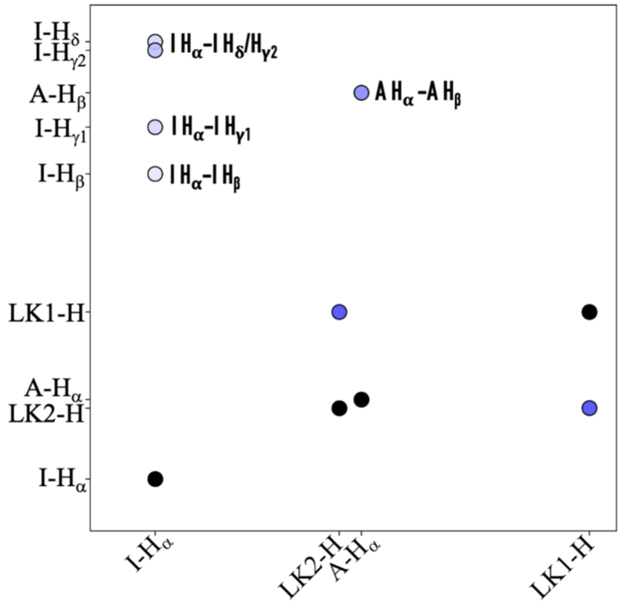


Figure S11. [1]H-[1]H NOEs calculated from MD simulation of AIssIA at a high-pH, dilute state. Average $r^{-6}$ values are displayed by the intensity of blue color; diagonal peaks are shown as black circles.

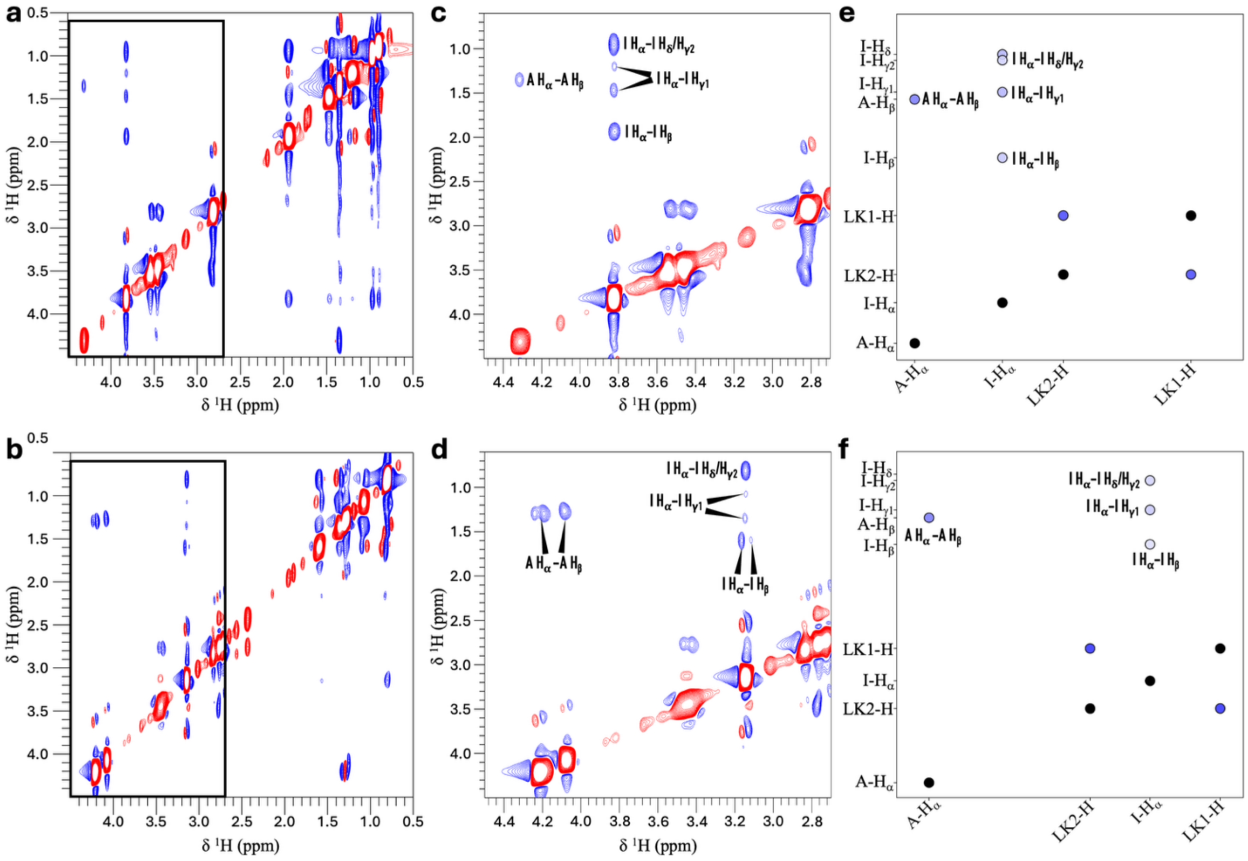


Figure S12. ¹H-¹H NOESY spectra of 50 mg/mL IAssAI. Full spectra are displayed in (a) for pH 2 and (b) for pH 13; enlarged view of the boxed regions in (a) and (b) are displayed in (c) and (d), respectively. ¹H-¹H NOEs calculated from MD simulations of IAssAI in the dilute state are displayed in (e) for low pH and (f) high pH.

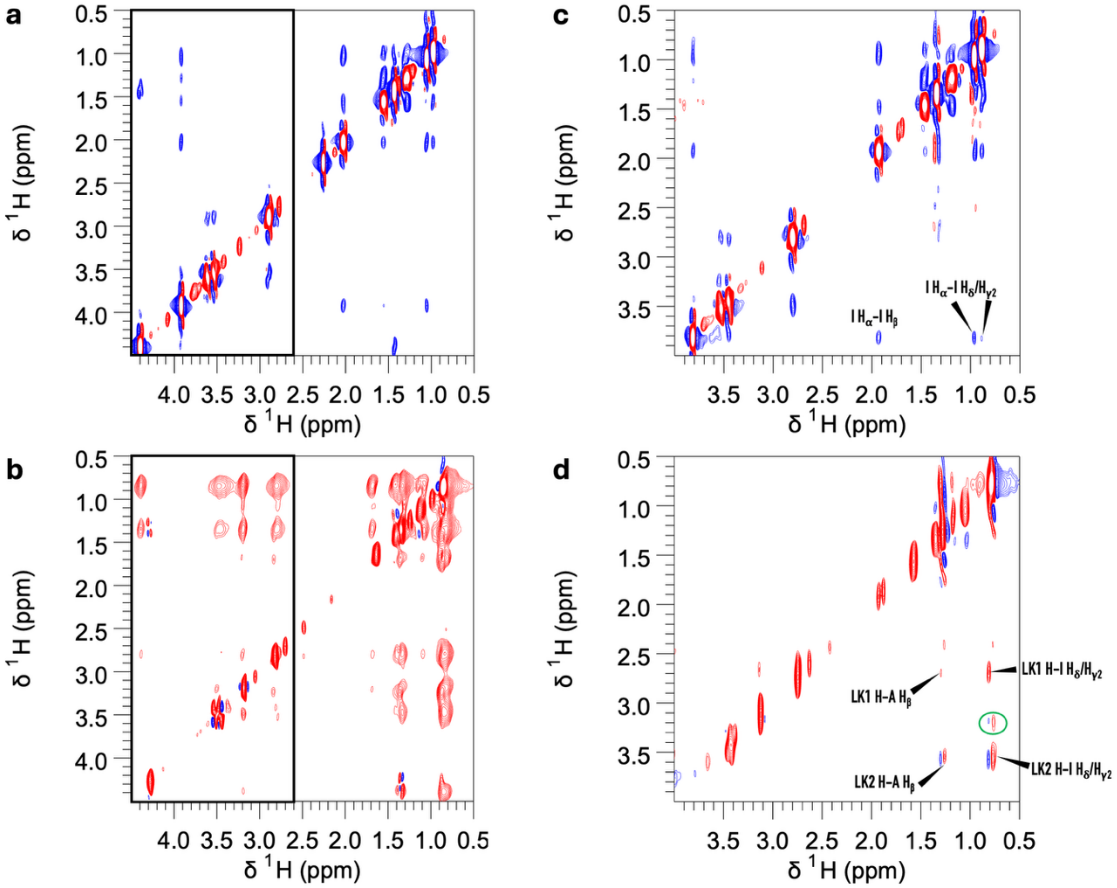


Figure S13. $^{1}H$-$^{1}H$ NOESY spectra of IAssAI, at 100 mg/mL and (a) pH 2 and (b) 13, or at 200 mg/mL and (c) pH 2 and (d) 13. Black rectangular boxes in (a) and (b) indicate regions that are shown in an enlarged view in Figure S14a, b. In (b) and (d), blue contours are artifacts arising from imperfect phasing.

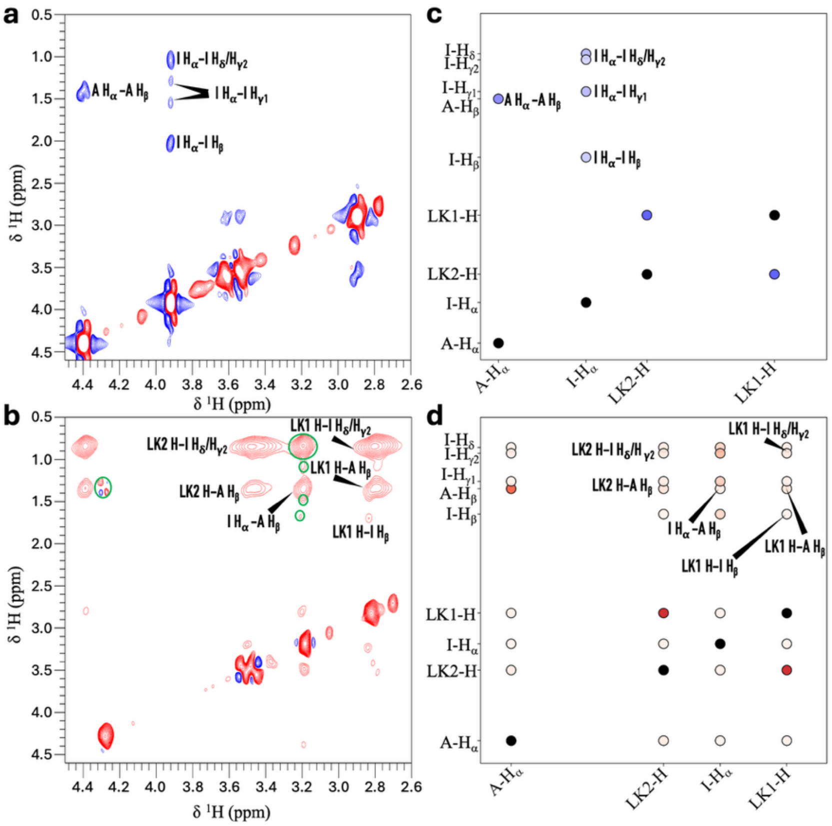


Figure S14. Enlarged view of Figure S13 and calculated $^{1}H$-$^{1}H$ NOEs from MD simulations. (a, b) Enlarged view of the boxed regions in Figure S13a, b. (c, d) $^{1}H$-$^{1}H$ NOEs calculated from MD simulations of IAssAI in the low-pH, dilute or high-pH, dense state.

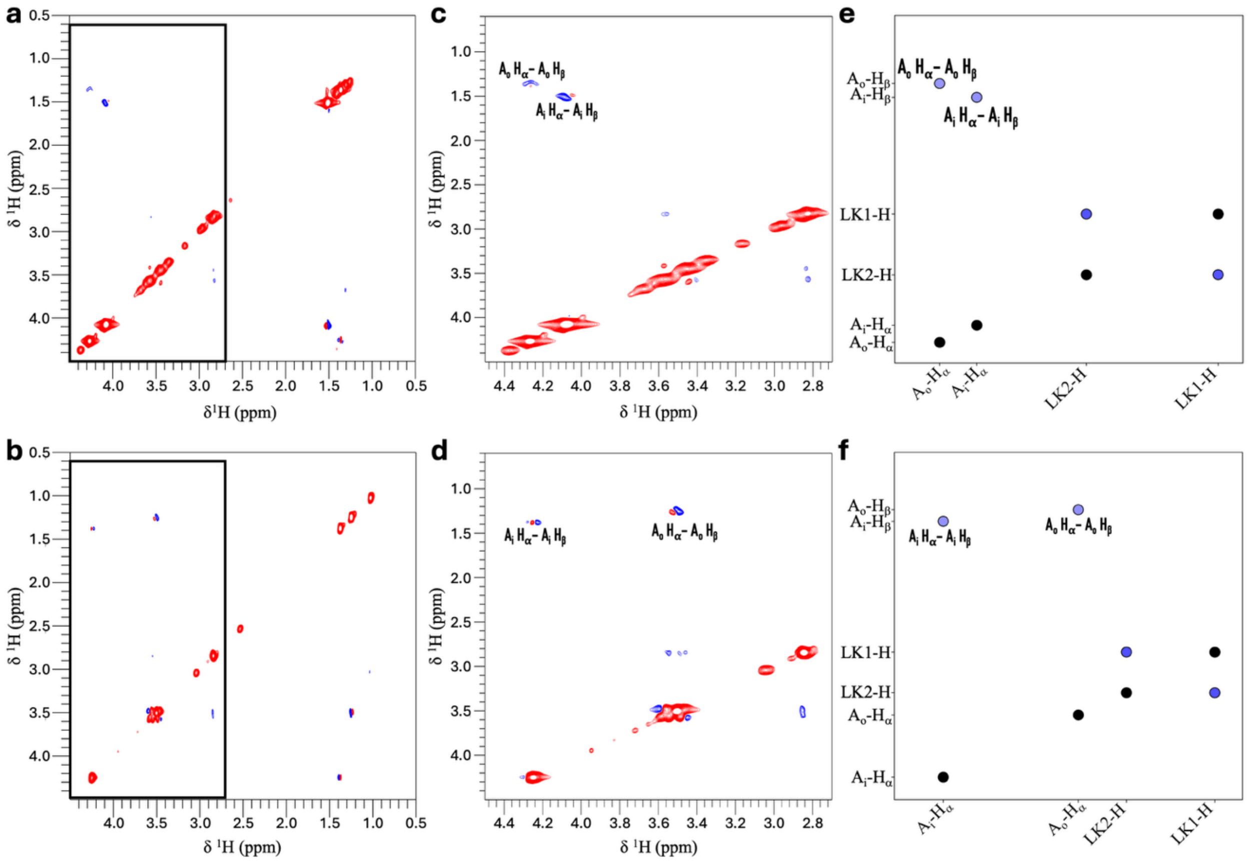


Figure S15. $^{1}$H-$^{1}$H NOESY spectra of 20 mg/mL AAssAAI. Full spectra are displayed in (a) for pH 2 and (b) for pH 13; enlarged view of the boxed regions in (a) and (b) are displayed in (c) and (d), respectively. $^{1}$H-$^{1}$H NOEs calculated from MD simulations of AAssAA in the dilute state are displayed in (e) for low pH and (f) high pH.

Videos

S1. Gel formation upon adding two drops of 5-M NaOH into 1 mL of 10-mg/mL AIssIA. Total time is 140 s.

S2. Conversion of gels into droplets after mixing AAssAA into AIssIA (5:1 molar ratio). For scale bar, see Figure 5c. The total time is 18 min.

S3. Conversion of gels into droplets after mixing AAssAA into AIssIA (5:1 molar ratio). For scale bar, see Figure S2b. The total time is 197.5 s.
